# The language network responds robustly to sentences across diverse tasks

**DOI:** 10.64898/2025.12.02.691902

**Authors:** Ruimin Gao, Chandler Cheung, Matthew Siegelman, Alvincé L. A. Pongos, Hope H. Kean, Alyx Tanner, Evelina Fedorenko, Anna A. Ivanova

## Abstract

A network of left frontal and temporal brain areas supports language comprehension and production, implementing computations related to word retrieval and combinatorial linguistic processing. Here, we ask: to what extent are responses to language in this language network stable across task contexts, and how does this stability compare to task sensitivity in the domain-general multiple demand (MD) network? Participants (n=52) read sentences and nonword lists under six task conditions, including passive reading, reading with a memory probe after each stimulus, and reading and answering questions that require deep semantic engagement. The sentences>nonwords contrast isolated the same set of language-responsive voxels across all tasks; the locations of those voxels were participant-specific, highlighting the value of individual-specific functional localization. We therefore conclude that language localization is robust to task variation. We then examined the magnitudes and fine-grained activation patterns in these language-responsive voxels (the language network) and in the domain-general MD network, to test whether task demands modulate linguistic computations and/or recruit a distinct brain system. The language network responded robustly to sentences across all tasks, with somewhat higher responses to semantically engaging tasks. In contrast, the MD network responded to both sentences and nonwords in the presence of a task, which warrants caution when using language paradigms that include task demands, as such paradigms engage two independent networks. A multivariate analysis further revealed that stimulus information is more easily decodable in the language network, whereas task information is more decodable in the MD network. These results suggest that the language and MD networks perform complementary functions during task-driven language comprehension, with the language network primarily extracting information from linguistic input and the MD network determining the appropriate response to the task.

## 1 Introduction

The human brain has a dedicated set of regions for processing language (Fedorenko et al., 2024). These regions, known as the language network, respond to a broad range of linguistic conditions across multiple levels of representation (from words to sentences and discourse): they are active during both production and comprehension, process written, spoken, and signed language, and respond to typologically diverse languages (Deniz et al., 2019; Fedorenko et al., 2010; Malik-Moraleda, Ayyash et al. 2022; Menenti et al., 2011; Neville et al., 1998; Regev et al., 2013; Scott et al., 2017; Silbert et al., 2014; Vagharchakian et al., 2012). However, the effects on the language responses of *task contexts* remain not well understood.

Language production almost always occurs in a context of a top-down goal—to express a particular meaning, either endogenously defined or driven by external demands (e.g., when answering a question). Language comprehension, on the other hand, may or may not be accompanied by top-down demands. Indeed, much evidence suggests that the language network is robustly engaged in passive, task-free comprehension paradigms (e.g., Bedny et al., 2011; Fedorenko et al., 2010; Lipkin et al., 2022; Malik-Moraleda, Ayyash et al. 2022; Scott et al., 2017), and when listening to naturalistic narratives (Blank & Fedorenko, 2017; Regev et al., 2013; Shain et al., 2022; Sueoka et al., 2024; Lerner et al., 2011; Wilson et al. 2008). This observation aligns with the fact that humans often extract meaning from language spontaneously, such as when overhearing a conversation or reading ads. But some evidence indicates that attention can affect responses in the language network, both positively and negatively. If a person is keenly interested in the topic of the text, their language areas respond more strongly (Olson et al., 2024). On the flip side, when participants engage in a challenging perceptual task that diverts their attention from linguistic content, activity in the language network declines substantially (Cohen et al., 2021; Ivanova et al., 2021).

Understanding the extent to which task demands modulate responses in the language network is important both theoretically and practically. Theoretically, this knowledge can help understand where the language network falls in the cortical hierarchy. In particular, brain regions vary widely in the extent to which their activity patterns are modulated by task context. Low-level sensory regions, such as the primary visual and auditory cortices, largely respond to the same stimulus in the same way regardless of task (Elhilali et al., 2007; Kay et al., 2008); their activity is only weakly modulated by attention (Jäncke et al., 1999; Poghosyan & Ioannides, 2008; Somers et al., 1999), and the auditory cortex responds to sounds even during sleep (Issa & Wang, 2008; Sela et al., 2016). Higher-level perceptual regions, such as the inferotemporal visual cortex, are more sensitive to task demands (Harel et al., 2014; Koida & Komatsu, 2007; McKee et al., 2014; Sigala & Logothetis, 2002) but are still primarily driven by the stimulus inputs (McMains & Kastner, 2011, Bugatus et al., 2017, Keller et al., 2022) and show robust engagement even when participants are focusing on unrelated tasks (Marvi et al., 2025). In contrast, some brain regions are strongly task-driven: for example, the fronto-parietal multiple demand (MD) network, implicated in goal-directed action and some aspects of reasoning, responds only during challenging cognitive tasks across diverse stimuli and paradigms (Assem et al., 2020; Duncan & Owen, 2000; Zanto & Gazzaley, 2013). Does the language network resemble perceptual regions, which show relatively stable responses across task contexts, or is it more similar to the MD network, which shows strong modulation by task demands?

On the practical side, a common way to identify the language network is with a passive comprehension contrast, such as simply reading sentences vs. nonword sequences (Fedorenko et al., 2010) or listening to sentences/short passages vs. to incomprehensible speech, such as speech in an unfamiliar language, speech played backwards, or acoustically degraded speech (e.g., Malik-Moraleda, Ayyash et al. 2022; Olson et al., 2023). The passive nature of the task makes it challenging to assess the level of stimulus engagement or the degree of comprehension in the absence of an explicit behavioral measure. Neural response magnitude can provide only an indirect indication of processing and does not always reliably index comprehension per se (e.g., Bautista & Wilson, 2016). These arguments are often used to defend a choice to include an explicit task in language paradigms. However,an explicit task may engage brain regions that support non-language-specific computations, such as general attention and working memory demands. Indeed, a meta-analysis of 30 fMRI language paradigms by Diachek et al. (2020) revealed that the language network responds similarly strongly, on average, to passive comprehension conditions and to conditions with active tasks, but the latter paradigms additionally engage the MD network, which shows no response during passive comprehension. Given that the language and the MD networks have been established to be functionally distinct (Braga et al., 2020; Du et al., 2024; Fedorenko & Blank, 2020; Mineroff, Blank et al., 2018; Quillen et al., 2021), the use of language paradigms that include an explicit task may be undesirable, because the elicited brain responses become difficult to interpret with respect to the cognitive processes they reflect (for additional discussion, see Ozernov-Palchik, O’Brien et al., in press; Billot, Jhingan et al., 2025).

One challenge with past comparisons of language paradigms with vs. without explicit tasks is that they have been indirect, performed across participants and across paradigms that differ in more than simply the presence or type of task, leaving some uncertainty as to how task demands affect brain responses to language. To fill this gap, we here systematically vary task demands in the same participants for the same stimuli across six tasks, while keeping other aspects of the paradigm as consistent as possible. The goal of this research is three-fold: i) to determine whether language localizers that use different tasks produce comparable results; ii) to better understand the effects of tasks during sentence-level comprehension with respect to the engagement of the domain-general MD network, and more generally, iii) to understand the language and the MD networks’ contributions to sentence-level comprehension under different task conditions, and to clarify how the language network fits within the broader continuum of stimulus- vs. task-driven neural responsiveness.

To foreshadow our key findings: across all (six) tasks, the language network responded robustly to the sentences>nonwords contrast, but the sentence conditions with a task additionally engaged the MD network. The presence of a deeper, semantic task led to a stronger response to the sentence condition in the language regions, indicating task-related modulation of response magnitude; however, the network’s core functional selectivity (sentences > nonwords) was preserved across task contexts. In contrast, the activity in the MD network scaled strongly with cognitive demand across stimulus types.

## 2 Methods

### 2.1 Participants

52 individuals (mean age = 23.9 years, SD = 3.8; 31 female) from MIT and the surrounding community participated for payment. 42 participants were right-handed and one was left-handed (but with typical, left-lateralized language responses), as assessed by Oldfield’s (1971) handedness questionnaire; handedness information was not collected for the remaining 9 participants (see **Supplemental Figure 1** for the laterality index of the included participants). Six additional participants were tested, but excluded from the analyses: three due to excessive head movement, two due to drowsiness, and one due to right-lateralized language responses. All participants had normal or corrected-to-normal vision.

### 2.2 Design, materials, and procedure

Six different versions of the language localizer task were used in the current study (**Figure 1**). Each participant completed between 3 and 5 versions (with distinct sets of experimental materials; materials were rotated across task versions across participants), and each task version was completed by between 19 and 52 participants (**Table 1**). All task comparisons are performed within participants using mixed-effects models with participants as a random effect to appropriately handle data imbalance. Across all versions, the critical condition required participants to read 12-word-long sentences (e.g., KNOWLEDGE GAINED FROM STUDYING EARTHQUAKE WAVES HAS BEEN APPLIED IN VARIOUS FIELDS), and the control condition required them to read lists of unconnected, pronounceable nonwords (e.g., KNORNEPTS GOWLED TROR CHOVYING EERSHQUORT WORED HOR BOUN ODSTIED EN NONIOUS FIECED). The sentences (a total of 5 sets, n=48 sentences each) were drawn from a language corpus (Francis & Kučera, 1964); the nonword lists were created by turning the words comprising the sentences into nonwords using the Wuggy software (Keuleers & Brysbaert, 2010). The materials are available at https://osf.io/zyfmu.

**Figure 1.**
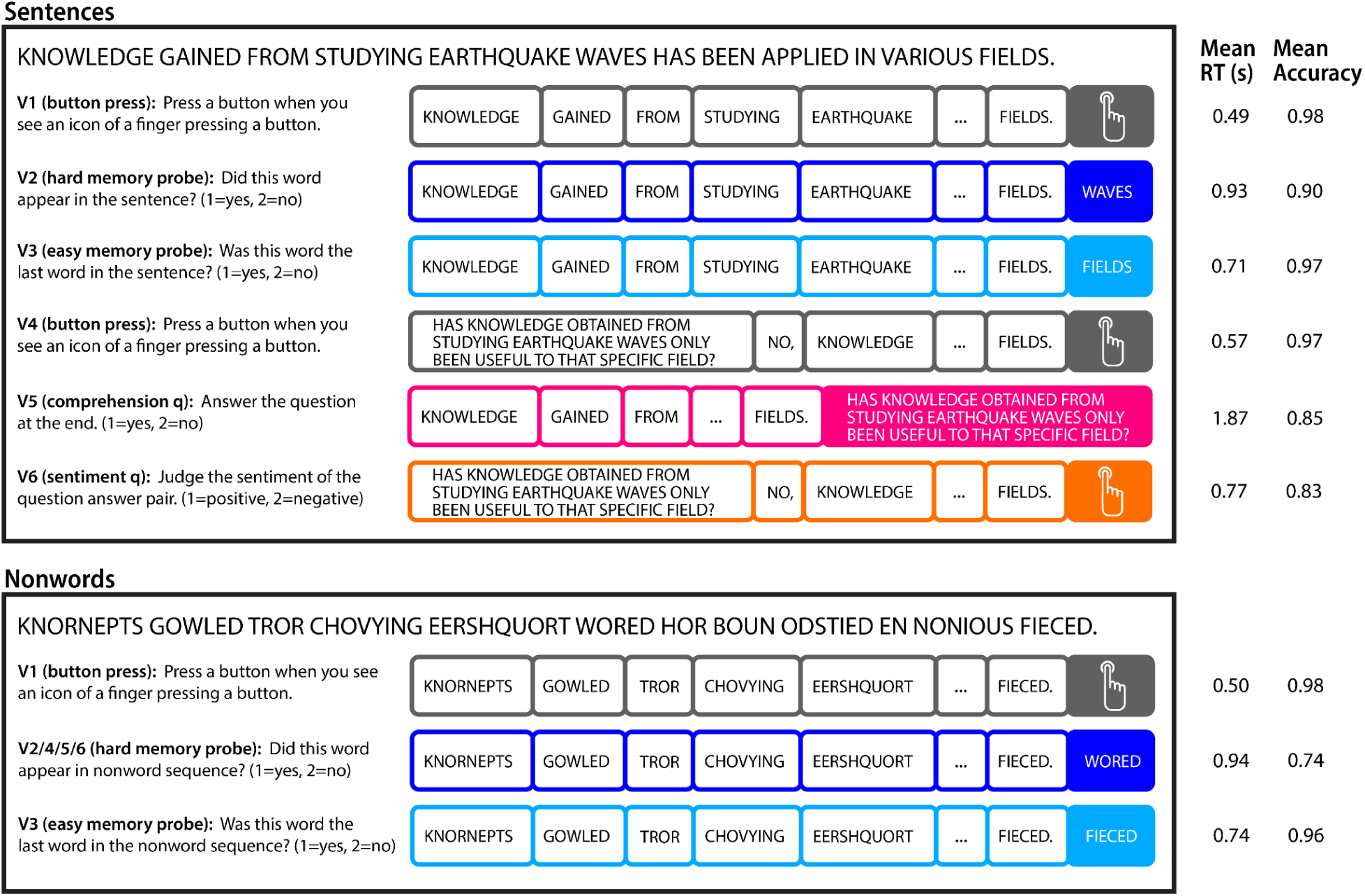
Design of the experiment. The sentences and nonword lists were presented in the center of the screen, one word/nonword at a time, at the rate of 400 ms per word/nonword. In V4-V6, where a question was included, the question was presented all at once for 2,600 ms. In V1 and V4, participants were simply asked to press a button when they saw the finger icon appear at the end of the sentence/nonword list. In V2 and V3, participants had to decide whether the probe word/nonword was present in the preceding sentence/nonword list; in V3, the correct probe was the same as the last word/nonword, which makes the task easy (effectively, deciding whether or not you just saw this word/nonword). For the sentence conditions, in V5, participants were asked to answer a yes/no comprehension question, and in V6, they were asked to make a sentiment judgment about the content of the question-answer pair. For V1-V3, the task was identical between sentences and nonword lists, for V4-V6, the hard memory probe version was used for the nonword lists. (See section 2.2 for details.)

**Table 1.** Participant counts for each unique combination of completed language task versions.

| Task | V1 | V2 | V3 | V4 | V5 | V6 | # Participants |
| --- | --- | --- | --- | --- | --- | --- | --- |
|  | X | X | X |  | X | X | 18 |
|  | X | X | X |  | X |  | 10 |
|  | X | X | X |  |  |  | 1 |
|  | X | X |  | X | X | X | 5 |
|  | X | X |  | X | X |  | 1 |
|  | X | X |  | X |  | X | 1 |
|  | X | X |  |  | X | X | 1 |
|  | X | X |  |  | X |  | 1 |
|  | X |  | X | X | X | X | 6 |
|  | X |  | X | X | X |  | 1 |
|  | X |  | X |  | X | X | 1 |
|  | X |  |  | X | X | X | 3 |
|  | X |  |  | X | X |  | 1 |
|  | X |  |  | X |  | X | 1 |
|  | X |  |  |  |  | X | 1 |
| **Total per task** | 52 | 38 | 37 | 19 | 48 | 37 | 52 |

In three versions (V1-V3), the sentence trials consisted of a single sentence presented one word at a time (**Figure 1**).These versions have been commonly used in past work (e.g., Fedorenko et al., 2010; Hu, Small et al., 2023; Lipkin et al., 2022; Malik-Moraleda, Ayyash et al. 2022). In ***Version 1*** (V1, “button press”), each sentence/nonword list was followed by a screen showing a picture of a finger pressing a button and participants were asked to press a button whenever they saw this picture (for the timing details, see **Supplemental Table 1**). In Versions 2 and 3, each sentence/nonword list was followed by a memory probe word/nonword and participants had to press a button to decide whether the probe was present in the preceding stimulus (button 1 for yes, button 2 for no). In ***Version 2*** (V2, “hard memory probe”), the correct word/nonword probe could come from any position in the sentence/nonword list; in ***Version 3*** (V3, “easy memory probe”), the correct probe came from the last position, which effectively turns this task into a decision about whether the probe was the last word/nonword.

In the sentence condition in the remaining three versions, participants read question-answer pairs. ***Version 4*** (V4, “button press”) was similar to V1, except that the sentence was preceded by a comprehension question presented on the screen all at once. The sentence was then presented (one word at a time, as in V1-V3, but with “Yes,” or “No,” appended to the beginning), such that the sentence provided the answer to the question. This version was designed as a control for Versions 5 and 6 that matches the amount of language material in the sentence condition. Versions 5 and 6 were designed to encourage deep engagement with the content of the sentences. In ***Version 5*** (V5, “comprehension q(uestion)”), each sentence was followed by a yes/no comprehension question (shown on the screen all at once, as in V4, except *after* the sentence), and participants were asked to answer it (button 1 for yes, button 2 for no). In ***Version 6*** (V6, “sentiment q(uestion)”), the sentence trials were structured as in V4, but participants were instead asked to make a judgment about the emotional valence of the question-answer pair (button 1 for positive content, button 2 for negative content). Participants were told that some sentences might be only a little bit positive or negative, or even seem totally neutral; in these cases, they were asked to make their best judgment. The nonword condition in V4-V6 used the hard memory probe task from V2.

The button press versions (V1 and V4) were designed to be as close as possible to passive comprehension with a very simple task that has nothing to do with the stimulus and simply serves to ensure that participants are awake; the button press also helps match these conditions to the other conditions for the motor demands. The remaining tasks—referred to as ‘active tasks’ in the remainder of the manuscript—all relate to the stimulus, and require greater attention to the stimulus and additional cognitive effort.

The sentences and nonword lists were presented in the center of the screen, one word/nonword at a time, at the rate of 400 ms per word/nonword. In Versions 4-6, the comprehension question was displayed all at once for 2,600 ms before (in V4 and V6) or after (in V5) the sentence. Each trial was preceded by a 100 ms blank screen. A blocked design was used with 3 sentence/nonword-list trials per block. Each run consisted of 16 experimental blocks (8 per condition), with fixation blocks (14 s each included at the beginning of the run after every four experimental blocks). The order of sentence and nonword blocks was palindromic within the run and counterbalanced across runs. Across versions, run durations varied between 5 min 58 sec and 8 min 22 sec. For each version, a given participant completed two runs.

In addition to the critical language tasks, the majority of the participants (45 of the 52) completed a spatial working memory task, which was used to localize the Multiple Demand network (Assem et al., 2020; Blank et al., 2014; Duncan, 2010; Mineroff, Blank et al., 2018). Briefly, participants viewed sequences of spatial locations flash up within a 3 × 4 grid, one at a time for a total of 4 locations (in the easy condition) or two at a time for a total of 8 locations (in the hard condition). At the end of each trial, they were shown two sets of locations and had to choose the set they just saw (see **Supplementary Figure 3** for details).

### 2.3 fMRI data acquisition

Structural and functional data were collected on the whole-body, 3 Tesla, Siemens Trio scanner with a 32-channel head coil, at the Athinoula A. Martinos Imaging Center at the McGovern Institute for Brain Research at MIT. T1-weighted structural images were collected in 176 sagittal slices with 1 mm isotropic voxels (TR = 2,530 ms, TE = 3.48 ms). Functional, blood oxygenation level dependent (BOLD), data were acquired using an EPI sequence (with a 90° flip angle and using GRAPPA with an acceleration factor of 2), with the following acquisition parameters: thirty-one 4 mm thick near-axial slices acquired in the interleaved order (with 10% distance factor), 2.1 mm x 2.1 mm in-plane resolution, FoV in the phase encoding (A >> P) direction 200 mm and matrix size 96 mm x 96 mm, TR = 2,000 ms and TE = 30 ms. The first 10 s of each run were excluded to allow for steady state magnetization.

### 2.4 Behavioral performance in the scanner

We analyzed response times using a linear mixed-effects model and accuracy using a binomial mixed-effects model. Both models included fixed effects of stimulus type (sentence vs. nonword list), linguistic demand (question–answer pair vs. single sentence), task (button press, hard memory probe, easy memory probe, comprehension question, sentiment question), and the task × stimulus-type interaction. For both models, we included by-participant random intercepts and random slopes for task and stimulus type. For the response time model, we additionally included a random intercept for the stimulus set, which significantly improved model fit (p < .05). For the accuracy model, including the stimulus set did not improve model fit (p = 0.621), so it was omitted.

To evaluate the accuracy of in-scanner sentiment ratings, we collected responses from 100 human participants on Amazon Mechanical Turk. After filtering for accuracy on attention check questions, native language (English), and native country (United States), we retained data from 80 participants. Each participant rated 1/4 of the sentences, leading to an average of 19.96 responses per sentence.

Runs with accuracies that were below 63.1%, which is a value that falls three standard deviations below the overall mean (across participants, versions, and conditions) were excluded (5 of the 472 runs), except for cases where the buttons were swapped or responses were not recorded due to equipment malfunction. In addition, we ensured that each participant had two runs per version, so if one run within a version had below-threshold accuracy, we excluded the version altogether for that participant (this criterion led to the exclusion of 5 additional runs).

### 2.5 fMRI data preprocessing

fMRI data were analyzed using SPM12 (release 7487), CONN EvLab module (release 19b), and other custom MATLAB scripts. Each participant’s functional and structural data were converted from DICOM to NIFTI format. All functional scans were coregistered and resampled using B-spline interpolation to the first scan of the first session (Friston et al., 1995). Potential outlier scans were identified from the resulting subject-motion estimates, as well as from BOLD signal indicators, using default thresholds in CONN preprocessing pipeline (5 standard deviations above the mean in global BOLD signal change, or framewise displacement values above 0.9 mm; Nieto 2020), and used as regressors of no interest in first-level analyses. Functional and structural data were independently normalized into a common space [the Montreal Neurological Institute (MNI) template; IXI549Space] using SPM12 unified segmentation and normalization procedure (Ashburner & Friston, 2005) with a reference functional image computed as the mean functional data after realignment across all timepoints omitting outlier scans. The output data were resampled to a common bounding box between MNI-space coordinates (−90, −126, −72) and (90, 90, 108), using 2 mm isotropic voxels and fourth-order spline interpolation for the functional data, and 1-mm isotropic voxels and trilinear interpolation for the structural data. Last, the functional data were smoothed spatially using spatial convolution with a 4 mm FWHM Gaussian kernel. For all experiments, effects were estimated using a general linear model (GLM) in which each experimental condition was modeled with a boxcar function convolved with the canonical hemodynamic response function (HRF) (fixation was modeled implicitly, such that all timepoints that did not correspond to one of the conditions were assumed to correspond to a fixation period). Temporal autocorrelations in the BOLD signal time series were accounted for by a combination of high-pass filtering with a 128 s cutoff, and whitening using an AR (0.2) model (first-order autoregressive model linearized around the coefficient a = 0.2) to approximate the observed covariance of the functional data in the context of restricted maximum likelihood estimation. In addition to experimental condition effects, the GLM design included first-order temporal derivatives for each condition (included to model variability in the HRF delays), as well as nuisance regressors to control for the effect of slow linear drifts, subject-motion parameters, and potential outlier scans on the BOLD signal. For the univariate analyses, we modeled the data using condition-level regressors; for the multivariate decoding analyses, we extracted estimates for the individual blocks.

### 2.6 Defining language and MD functional regions of interest (fROIs)

Language and MD fROIs were defined using the group-constrained subject-specific (GSS) approach (Fedorenko et al., 2010; Julian et al., 2012) where a set of spatial masks, or parcels, is combined with each individual participant’s localizer activation map, to constrain the definition of individual fROIs. The parcels delineate the expected gross locations of activations for a given contrast based on prior work and large numbers of participants, and are sufficiently large to encompass the inter-individual variability in the locations of functional areas. For the language fROIs, we used a set of ten parcels derived from a group-level probabilistic activation overlap map for the sentences > nonwords contrast in 220 participants and used in much prior work (e.g., Malik-Moraleda, Ayyash et al. 2022; Olson et al., 2024; Schrimpf et al., 2021; Shain et al., 2022; Tuckute et al., 2024). For each hemisphere, the parcels included two regions in the inferior frontal gyrus (IFG, IFGorb), one in the middle frontal gyrus (MFG), and two in the temporal lobe (AntTemp and PostTemp). The angular gyrus regions, which were included in the original set of areas reported in Fedorenko et al. (2010), were not included because of their systematically distinct response profile relative to other language-network regions and low functional connectivity with the rest of the language network (e.g., Blank et al., 2014; Blank et al., 2016; Ivanova et al., 2019; Jouravlev et al., 2019; see Shain, Paunov, Chen et al., 2022 for discussion).. For the MD fROIs, we used a set of 20 parcels (10 in each hemisphere) derived from a group-level probabilistic activation overlap map for the hard > easy spatial working memory contrast in 197 participants and also used in many prior studies (e.g., Assem et al., 2020; Shain et al., 2022; Kean et al., 2025). The parcels included regions in the frontal and parietal lobes, as well as a region in the anterior cingulate cortex. The language and the MD parcels are available at https://www.evlab.mit.edu/resources-all/download-parcels. Within each parcel, we selected the 10% of voxels with the highest t-values for the contrast of interest (sentences > nonwords for the critical language tasks, hard > easy spatial working memory for the MD localizer).

For exploratory analyses, we defined functional regions of interest in another high-level cognitive network—the default mode network (DMN) (Buckner & DiNicola, 2019)—using a similar procedure. For these fROIs, we used a set of 10 parcels (5 in each hemisphere) derived from a group-level probabilistic activation map for the easy > hard spatial working memory contrast in 197 participants. The parcels included regions in the posterior cingulate cortex, in the medial frontal cortex, in the precuneus, and in the temporoparietal junction. The results are reported in **Supplementary Tables 6** and **9**, for completeness.

### 2.7 Response magnitude analyses

We evaluated the responses of the language and MD fROIs to the conditions of the critical language tasks. For each condition, the effect was averaged across the voxels within each fROI to derive a single value per fROI per participant. When estimating the responses to the conditions of the task that was used as the localizer for that fROI, the effects were estimated using across-runs cross-validation to ensure independence (Kriegeskorte et al., 2009).

### 2.8 Spatial overlap analyses

To assess the degree of overlap between language fROIs defined using different localizer task versions, we used the Dice coefficient measure (Dice, 1945). To equalize the amount of data between the within-task comparisons (where each data half consists of a single run) vs. across-task comparisons (where each task consists of two runs), for the latter, we evaluated the overlap between fROIs defined by localizer task A (run 1) and localizer task B (run 2), and vice versa, averaging across the splits to derive a single value per fROI per participant. In the analyses, we compared the within-participant between-task consistency to the within-task between-participant consistency.

### 2.9 Spatial correlation analyses

To investigate the activation patterns of voxel-level responses within each language fROI, we extracted vectors with that fROI’s voxelwise responses to a single stimulus type (sentences/nonwords) and a single language task version (2 stimulus types x 6 task versions = 12 conditions in total). We then computed Pearson correlation values for all pairwise condition combinations for each fROI. To ensure the independence of data used to estimate the voxelwise responses in each condition, we used separate runs to calculate each of the two effects that we compare (and, when possible, averaging between estimates for (a) Run 1 - Condition 1 vs. Run 2 - Condition 2 and (b) Run 1 - Condition 2 vs. Run 2 - Condition 1). Because we have a total of 2 runs per task version, when we use one run as the fROI localizer, we do not have enough remaining data to compare two effects from the same task version. As a result, we do not provide values for V1 Sentence/Nonword - V1 Sentence/Nonword correlation when using V1 as the localizer. The resulting Pearson correlation values were Fisher z-transformed for regression analyses.

### 2.10 Multivariate decoding of stimulus vs. task information

To investigate whether different stimulus types and different tasks evoke distinct, linearly separable response patterns in the language and the MD networks, we trained within-participant linear classifiers on block-level voxelwise responses to each condition. For this analysis, we only included V1, V2, and V3, because the task was identical between the sentence and nonwords conditions for these versions. To ensure that the results were not biased by the localizer version used to define the fROIs, we used V4 as the localizer. A total of 29 participants completed all V1, V2, and V3 tasks; of these, 21 also completed V4 and the MD localizer. These 21 participants were included in this analysis.

For each block and each fROI, we z-scored voxel responses across voxels (mean-centered and scaled to unit variance) to remove univariate differences between conditions.

We adopted five-fold cross-validation across blocks to split the data into train and test sets. To ensure generalizability, we evaluated the performance of four classifiers—a correlation-based classifier, k-nearest neighbors (kNN), logistic regression, and support vector machines (SVM). The results were similar across these approaches, so we report the correlation-based classifier results in the main text, and the rest of the results in **Supplemental Table 12**. The decoding analyses were conducted using the pyMVPA package (Hanke et al., 2009).

### 2.11 Statistical analyses

We conducted the analyses using mixed-effects models implemented in the R package lmerTest (Kuznetsova et al., 2017). Separate models were fit for the left and right hemisphere language and MD networks.

#### Response magnitude analyses

We fit a mixed-effects model predicting fMRI activation (BOLD signal change) with fixed effects of linguistic demand (which varied across task versions (single sentence (used in V1-V3, and question-answer pair (used in V4-V6)), the interaction between the localizer task and linguistic demand to test whether responses to the sentence condition are modulated by task demand, and a three-way interaction between the localizer task, linguistic demand, and stimulus type (sentences, nonwords) to test whether nonword responses differ significantly from sentence responses for each localizer task. We included random intercepts for fROIs and participants, and random slopes for condition (sentences, nonwords) and task version where possible. For language-network analyses, we additionally included a random intercept for localizer-task version. When comparing the results across localizer versions used to define the fROIs, we fit separate models for each localizer version; and when comparing the results across fROIs, we fit separate models for each fROI; in both cases, an FDR correction was used to account for multiple comparisons.

#### Spatial overlap analyses

We fit a mixed-effects model predicting spatial overlap with fixed effects of comparison type (within-participant vs. between-participant) and task (within-task vs. between-task), and their interaction. Random intercepts were included for participant pairs and fROIs. Post-hoc comparisons with Tukey’s correction were conducted to examine: (1) within-participant within-task vs. within-participant between-task overlap for each task version, and (2) within-participant between-task vs. between-participant within-task overlap for each task version.

#### Spatial correlation analyses

We fit a mixed-effects model predicting Fisher-z-transformed correlation values with fixed effects of effect pair (sentences vs. sentences, sentences vs. nonwords, nonwords vs. nonwords), the interaction between linguistic demand (single sentence, question-answer pair), and the interaction between effect pair and task (simple button press (V1, V4), active task (V2-V3, and V5-V6)). Random intercepts were included for participants and fROIs in all models; for language-network analyses, we additionally included a random intercept for localizer-task version.

#### Decoding analyses

To test the significance of decoding accuracy, we conducted permutation tests (N = 1,000) to determine whether participants’ average decoding accuracies exceeded chance with an FDR correction. To compare decoding accuracy across networks, we fit a binomial mixed-effects model for prediction accuracy that included a fixed three-way interaction among training type (within- vs. across-task for stimulus type decoding, or within vs. across-stimulus-type for task decoding), network (LH language network vs. MD network), and testing group (task version (V1/V2/V3) for stimulus-type decoding or condition (sentences/nonwords) for task decoding). Random intercepts were included for participants and classifier type (correlation-based/kNN/logistic/SVM). Post-hoc comparisons with a Bonferroni correction were conducted to examine accuracy differences across networks for each combination of training type and testing group.

## 3 Results

### 3.1 All six task versions of the language localizer are effective at identifying the language areas within individual brains

First, we examined the responses to sentences and nonwords in the language network defined using different task versions of the localizer, and tested whether these responses were spatially consistent across fROIs (**Figure 2**). For all language task versions, the left-hemisphere language network showed a significantly stronger response to sentences compared to nonwords (ps < .001), regardless of the localizer used for fROI definition (see **Supplemental Table 2** for the statistical results). These effects also held for all five left-hemisphere language fROIs individually (ps < .001; **Supplemental Table 3**).

**Figure 2.**
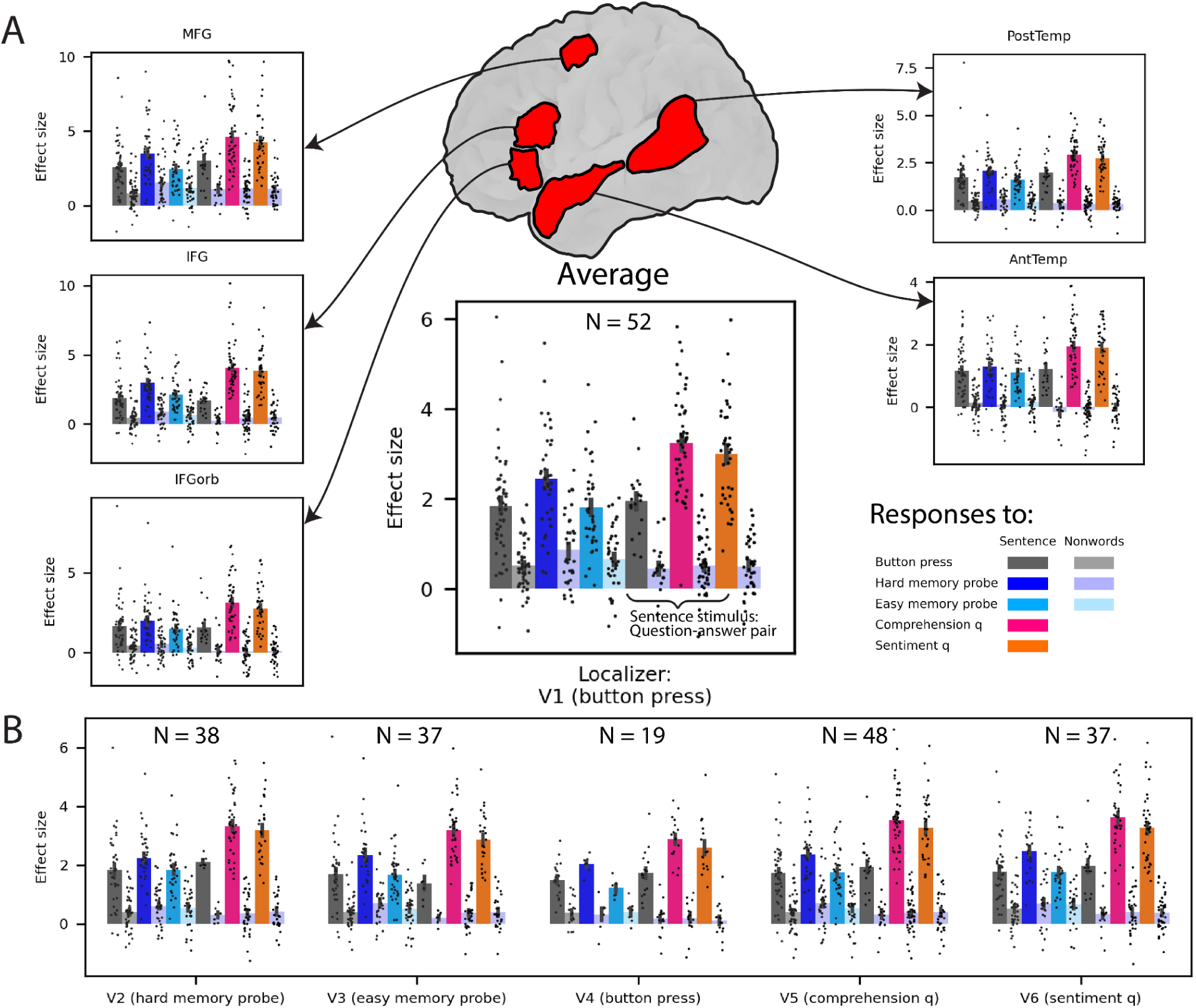
Responses of the core left-hemisphere language network to the sentence and nonword conditions in the six task versions. (A) Average network-level response profile (large bar graph) and fROI-level profiles (smaller bar graphs). The language fROIs were defined with language localizer V1 (button press), which every participant performed. Error bars denote the standard errors of the mean by participant; dots correspond to individual participants. (B) Average network-level response profiles for language fROIs defined using the other five localizer versions (V2-V6).

Next, we assessed the topographic consistency of the language fROIs defined using the same or different tasks within individuals, or—for comparison—using the same task across individuals. Within-participant overlap values were consistently high, both within each task (0.59 on average; **Figure 3**) and across tasks (0.50 on average). These values fall in the fair-to-good range according to commonly used benchmarks for interpreting overlap coefficients (Cicchetti, 1994). The within-task patterns did show significantly greater similarity compared to the between-task patterns, which suggests that different task variants activate slightly different sets of voxels (ps < .001, **Supplemental Table 10**). However, even the within-individual between-task values were substantially higher than the between-individual within-task values (0.13 on average, considered poor according to Cicchetti’s (1994) criteria) (ps < .001), consistent with earlier findings of substantial inter-individual variability in the precise locations of the language areas (Braga et al., 2020; Du et al., 2024; Fedorenko et al., 2010; Mahowald & Fedorenko, 2016; Shain & Fedorenko, 2025). These patterns were consistent across all five left-hemisphere language fROIs (**Supplemental Table 11**).

**Figure 3.**
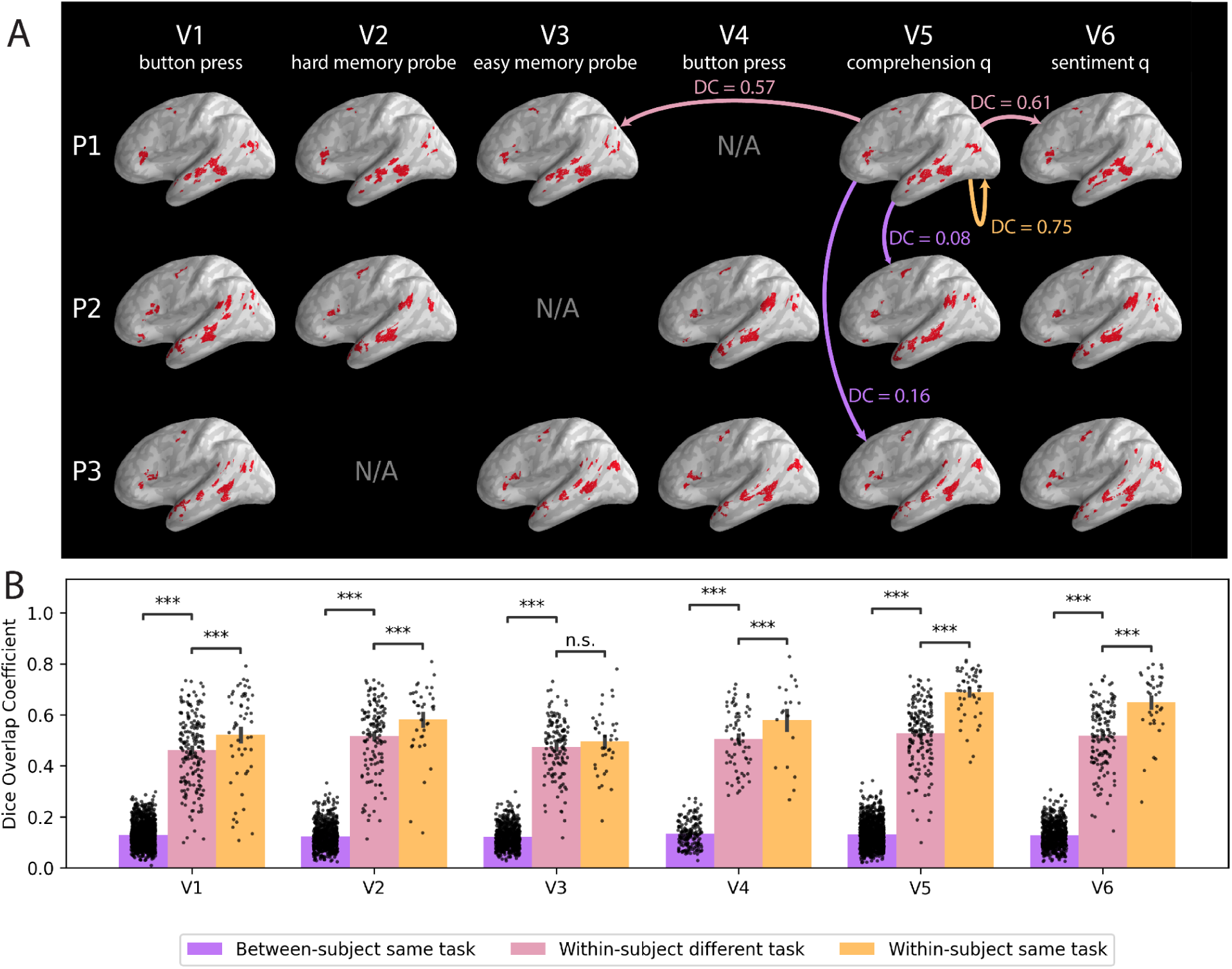
Language fROIs defined by different language localizer task versions. (A) Language fROIs defined in three sample participants (P1-P3). Each participant completed 5 of the 6 localizer versions. DC = Dice overlap coefficient. The Dice coefficient is calculated for the left-hemisphere language fROIs within participants within and between tasks (orange and pink arrows) or between participants for the same task (purple arrows) (<u>Methods</u>). (B) Average Dice overlap of left-hemisphere language fROIs within participants (orange and pink bars) and between participants for the same task (purple bar) (ns: not significant, *: p < .05, **: p < .01, ***: p < .001). Separate runs are used to define each of the two fROIs being compared (averaging between estimates for (a) Task 1 - Run 1 vs. Task 2 - Run 2 and (b) Task 1 - Run 2 vs. Task 2 - Run 1). Error bars denote the standard errors of the mean over participants / participant pairs; dots correspond to participants (for the within-participant comparisons) and to pairs of participants (for the between-participant comparison).

### 3.2 Task demands lead to higher responses in the language network but also engage the multiple demand network

The language and the multiple demand network were differentially affected by stimulus type vs. task in both univariate responses and activation patterns.

We investigated the effects of stimulus type and task on responses in the left and right hemisphere components of the language and MD networks. The two networks showed distinct activation profiles (**Figure 4A**). In the language network, we observed a robust sentence > nonword contrast for all tasks in both hemispheres (ps < .001; **Supplemental Table 4**). In contrast, the MD network showed higher responses to nonwords than to sentences in both hemispheres for V1-V4 (ps < .05); for V5 (comprehension question) and V6 (sentiment question), the sentence and nonwords conditions did not reliably differ. An important thing to note is that all sentence conditions with an active task (i.e., tasks requiring more than a simple button press at the end of a stimulus) elicited an above-baseline response in the MD network (ps < .05), and in some cases, this response was higher than the response to the nonwords condition (see **Supplemental Table 5**). This means that depending on the control condition, sentence comprehension with an active task may engage not only the language areas but also the MD network.

**Figure 4.**
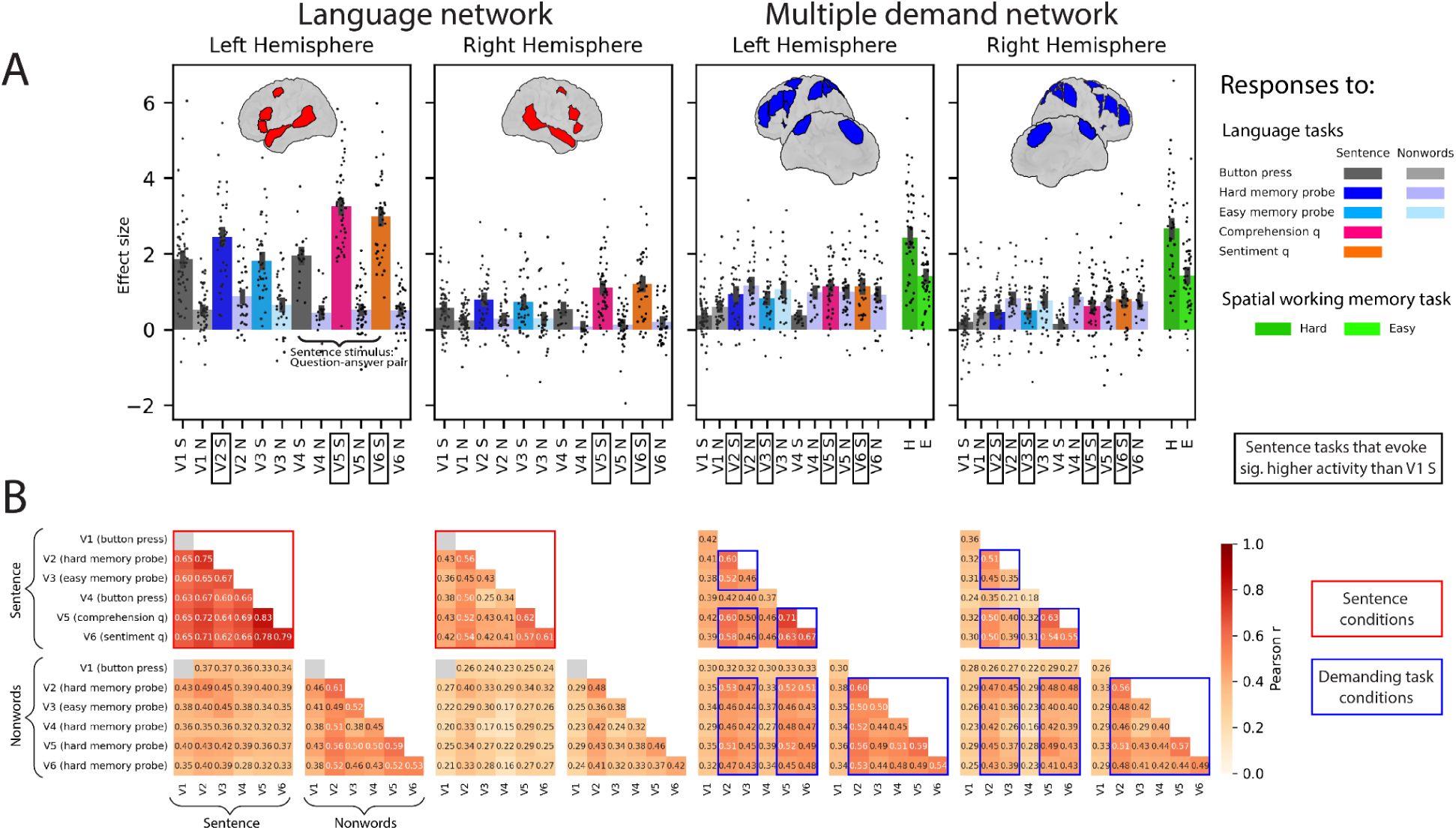
(A) Responses of the left-hemisphere and right-hemisphere language and MD networks to the sentence and nonword conditions in the six task versions. As in Figure 2, here and in the results in panel B, the language fROIs were defined with language localizer V1 (button press), which every participant performed. Error bars denote the standard errors of the mean by participant; dots correspond to individual participants. (B) Spatial correlation analyses of activation patterns within the networks. Separate runs are used to estimate each of the two effects being compared (averaging between the estimates for (a) Task 1 - Condition 1 vs. Task 2 - Condition 2 and (b) Task 1 - Condition 2 vs. Task 2 - Condition 1) .

Regarding task effects, the left-hemisphere (LH) language network showed increased activation for the sentence condition in V2 (hard memory probe) relative to V3 (easy memory probe) and relative to V1 (button-press) (ps < .001; **Supplemental Table 4**). However, this increase is ambiguous: it could reflect either direct task-induced modulation of language regions (e.g., attentional enhancement) or incidental task-induced re-processing of the linguistic input (e.g., subvocal rehearsal), which cannot be disentangled with the present design. No difference was observed for these contrasts in the RH language network, nor for the comparison of V3 (easy memory probe) relative to V1 for either the LH or the RH language network. Further, both the LH and RH language network showed increased activation for the sentence condition in V5 (comprehension question) and V6 (sentiment question) relative to the button-press version that was matched for the amount of linguistic input (V4), as well as relative to the sentence conditions in V1-V3 (all ps < .001). Importantly, however, these task-related effects were dwarfed by the effects of stimulus type. The average magnitude of response in the language network was not significantly predicted by the difficulty of that condition as reflected in average accuracies and response times (ps > .30 for both hemispheres; **Supplemental Figure 2**). For the sentence conditions, response magnitudes were significantly predicted by accuracy, with higher responses to more difficult conditions (i.e., conditions associated with lower accuracies) (ps < .005 for both hemispheres; **Supplemental Figure 3**) but not by response time (p = .109 for LH; p = .167 for RH). For the nonword conditions, response magnitudes were not significantly predicted by either difficult measure (all ps > .40).

In contrast, the MD network exhibited more robust task effects compared to the stimulus-type effects: in both hemispheres, for all active tasks, the sentence condition elicited a higher response than the corresponding button-press condition matched for the amount of linguistic content (i.e., V2 and V3 compared to V1; and V5 and V6 compared to V4) (all ps < .001; **Supplemental Table 5**). Furthermore, across the 12 conditions, the average response magnitude in the MD network was well predicted by the difficulty of that condition as reflected in average accuracies (ps < .05 for both LH and RH; **Supplemental Figure 2**) but not response times (p = .054 for LH and p = .173 for RH).).

To test whether fine-grained activation patterns are more similar for the same stimulus across tasks or for the same task across stimuli, we conducted a spatial correlation analysis of voxelwise responses. The multivariate patterns mirrored the univariate responses. Within both LH and RH language networks, the activation pattern was more consistent among the sentence conditions than among the nonword conditions (ps < .001; **Figure 4B**; **Supplemental Table 7**). However, for the MD network, this effect was observed only in the left hemisphere (p < .001; **Supplemental Table 8**). Furthermore, in both LH and RH language networks, the activation pattern for the sentence condition became more distinct from that for the nonword condition when the sentence conditions involved the additional linguistic demand of a question-answer pair (in V4-V6) (ps < .001). In contrast, in the MD network, the increased linguistic demand in the sentence conditions resulted in activation patterns that were more similar to those for the nonword conditions (ps < .001). Finally, both networks showed more consistent within-condition activation patterns under task conditions compared to passive reading (button press) versions matched for the amount of linguistic content (i.e., V2 and V3 vs.V1; and V5 and V6 vs. V4) (ps < .001). However, in the LH language network, this effect was substantially smaller than the sentences > nonwords effect (task effect: 0.129; sentences > nonwords effect: 0.362; the effects were more comparable in the RH language network: task effect: 0.136; sentences > nonwords effect: 0.149). In contrast, in the MD network, the task effect was substantially larger than the sentences > nonwords effect (LH: task effect: 0.281; sentences > nonwords effect: 0.112; RH: task effect: 0.255; sentences > nonwords effect: 0.016).

### 3.3 Fine-grained activity in the language network better distinguishes stimulus type but activity in the Multiple Demand network better distinguishes task type

Finally, to complement the pattern-similarity analyses, we asked whether stimulus (sentences vs. nonwords) and task (button press vs. hard memory probe vs. easy memory probe) could be explicitly decoded from the activation patterns within each network. We found that both stimulus and task could be reliably decoded in the LH language network and in the MD network, whether the classifier was trained and tested on the same task or trained on two tasks and tested on the left-out task (ps < .05; **Figure 5**; **Supplemental Table 12**). However, post-hoc tests revealed that, for both within- and between-task classification, the language network exhibited higher decoding accuracy for stimulus type compared to the MD network (average decoding accuracies 0.69 and 0.56, respectively; p < .001). In contrast, the MD network showed higher decoding accuracy for tasks than the language network, although the difference was less pronounced (0.42 vs. 0.40; p < .001, except when both training and testing were performed on the sentence condition: p = .072). Thus, although each network contains information about both stimulus type and task, they differ in how robustly they represent these two kinds of information, with stronger stimulus coding in the language network and stronger task coding in the MD network.

**Figure 5.**
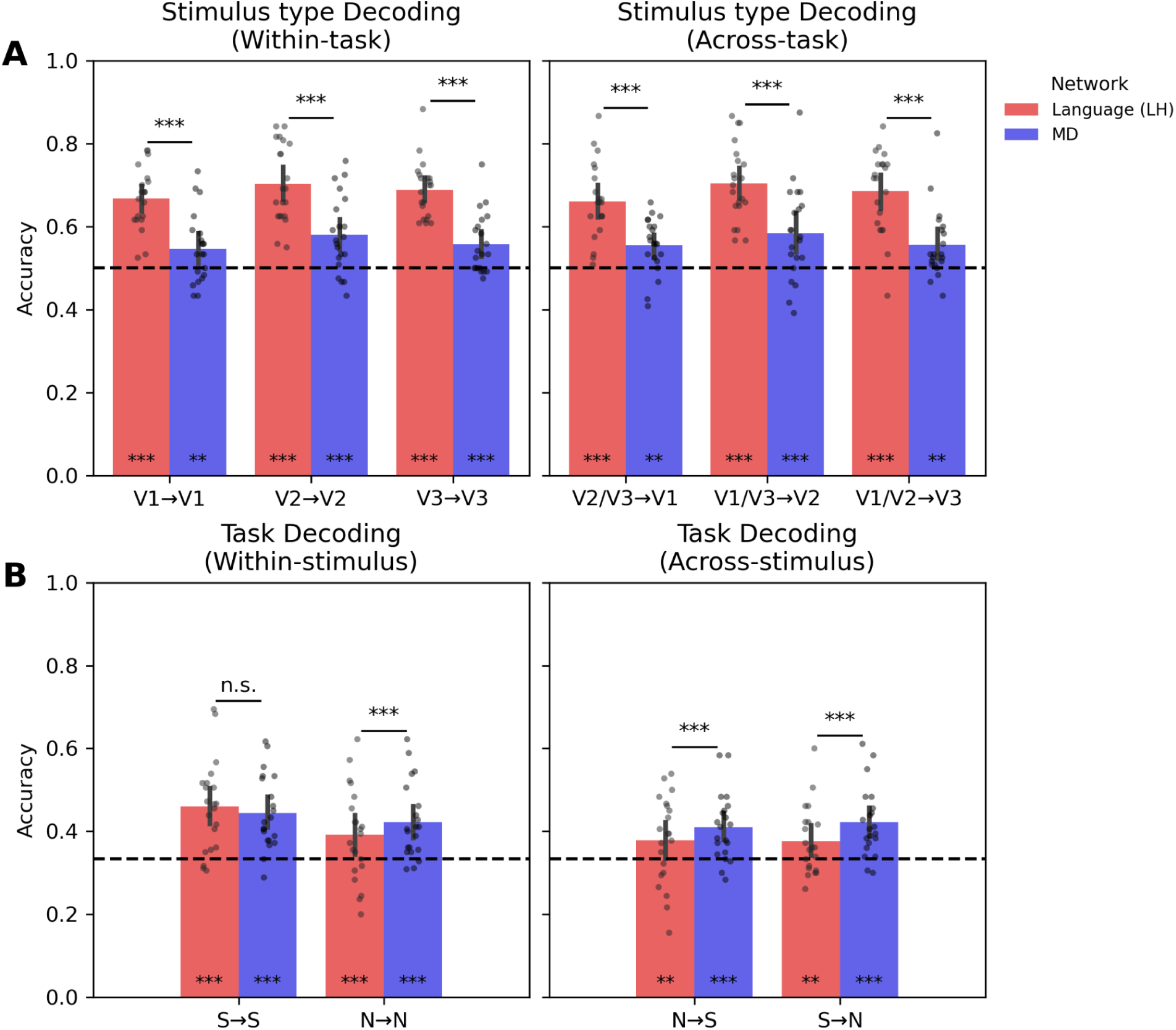
Multivariate pattern analysis results for the LH language network (red bars; see Supplemental Table 12 for the results of the RH language network) and the bilateral MD network (blue bars) in classifying stimulus (A) and task (B). (See Supplemental Table 12 for evidence that the results generalize across other types of classifiers.) Dotted lines indicate chance-level decoding accuracy: 0.5 for stimulus-type decoding (binary classification) and 1/3 for task decoding (three-way classification).

## 4 Discussion

In this study, we systematically varied task demands—from a simple button press to challenging memory probes and content-based comprehension questions/judgments—to test whether task contexts affect the localization and response profile of the language network in individual brains. We found that the language network localizer is robust to task variation, yielding a spatially consistent set of brain areas with remarkably similar activation profiles. More demanding tasks elicit somewhat stronger responses in the language areas, but they additionally engage a distinct non-language-selective network—the multiple demand (MD) network. Finally, the effects of stimulus type are stronger in the language than in the MD network, as reflected in both univariate responses and multivariate activation patterns, and the task effects are stronger in the MD network. Below, we discuss the theoretical and practical implications of these findings.

### 4.1 Responses to language in the language network are stable across task contexts

We found that the LH language network’s response during sentence comprehension remained robust across task contexts. Specifically, the language areas showed a consistently stronger response to sentences than to nonwords regardless of whether participants were simply engaged in attentive passive reading (in our button-press conditions), in tasks that required them to remember the words that composed the sentences (in the memory-probe conditions), or in tasks that required evaluating the semantic content of the sentence (in the comprehension-question and sentiment-judgment conditions). These results align with prior work demonstrating robust sentence > nonword effects across stimuli, presentation modalities, tasks, and languages (see Fedorenko et al., 2024 for review), and extend it by showing that these effects hold across six different task versions holding all other aspects of the design constant. Across the network and in each individual language region, the sentence conditions elicited a response that was several times higher than that elicited by the nonwords conditions. Moreover, the locations of these language-selective areas were highly similar across task versions within individuals (and much more similar than for the same task across individuals, in line with previously reported inter-individual topographic variability; Fedorenko et al., 2010; Braga et al., 2020; Shain & Fedorenko, 2025).

Although stimulus type (sentences vs. nonwords) explained the most variance in between-condition differences in the language network, we also detected moderate increases in activation during the more demanding tasks, such as those requiring memory retrieval or interpretive judgment (see e.g., Quillen et al., 2021 for earlier evidence of the language responses being modulated by the language task difficulty). This increase in the level of response for more demanding tasks is, however, consistent with at least two interpretations. One possibility is direct modulation of the language regions by task-related control or attentional signals. But another possibility is that task demands induce additional stimulus-related computations. In particular, to make a memory probe judgment or to answer a question about the content sentence, it may be necessary to rehearse or reactivate the sentence content, which would lead to additional activity in the language areas. These possibilities are difficult to distinguish empirically given that any increase in task demands is likely to lead to additional engagement with the stimulus. For example, the greater response observed for the hard relative to the easy memory probe can reflect increased engagement with the sentence content (e.g., maintenance or rehearsal), or sensitivity to domain-general task difficulty. However, other aspects of the results, discussed in Section 4.3, suggest that additional stimulus processing may be more likely as an explanation.

The patterns in the RH homotopic language regions largely mirrored those in the core LH regions: conditions that elicited stronger responses in the LH produced proportionally weaker responses in the RH. This pattern suggests a difference in response magnitude rather than a qualitative divergence in processing, consistent with prior functional localization work showing similar but attenuated responses in the RH (Fedorenko et al., 2010). Although some task-related increases observed in the LH (e.g. easy memory probe vs. button press) were not reliably present in the RH, this difference is parsimoniously explained by overall attenuation rather than distinct computations. Future work should directly test whether any conditions elicit genuinely different computations across hemispheres.

Taken together, these findings suggest a high degree of stability in the language network’s responses across task contexts. Whether similar stability extends to other modalities of comprehension or to language production remains to be established in future work.

### 4.2 Implications for functional localization of the language network

The significant variability in the precise locations of the language regions across individuals poses a problem for approaches that average data voxelwise across individuals (see Fedorenko, 2021 for discussion). One alternative is to define these regions functionally in each individual brain using a simple ‘localizer’ task (Saxe et al., 2006)—this approach has been standardly used in vision research for many years (e.g., Downing et al., 2001; Epstein & Kanwisher, 1998; Kanwisher et al., 1997) and has been extended to many other domains, including language (Fedorenko et al., 2010). The most common contrast used to identify the language areas is between language processing and a perceptually matched control condition (e.g., reading sentences vs. lists of nonwords; or listening to sentences/passages vs. to backwards speech), typically accompanied by a button press or a memory probe task to ensure engagement (Bedny et al., 2011; Fedorenko et al., 2010; Hiersche et al., 2024; Lipkin et al., 2022; Malik-Moraleda, Ayyash et al. 2022; Scott et al., 2017). Here, we investigated whether or how task demands affect the locations and functional profiles of the language regions when using sentences and nonword lists as the stimulus contrast.

Our findings show that within a given individual, the topography of the functional regions of interest (fROIs) defined by the reading-based sentences>nonwords contrast is highly consistent across six task versions—a testament to the robustness and generalizability of this localizer contrast. Within-participant overlap values of language fROIs were high both within each task, consistent with prior reports of topographic stability in the language network (e.g., Mahowald & Fedorenko, 2016), and importantly across tasks . In line with this topographic similarity, the response profiles of these fROIs were also similar, all showing a large and replicable across runs sentences>nonwords effect, and selectivity relative to a demanding non-linguistic task (a spatial working memory task). These results suggest that at least for neurotypical adults, researchers can use any of these versions of the localizer without compromising the validity or reproducibility of language network localization.

This robustness to task context adds to prior evidence of the robustness of this localizer contrast (between language processing and a perceptually similar control condition where the content is indecipherable) to modality of presentation (reading, listening, or audio-visual; Fedorenko et al., 2010; Olson et al., 2024; Scott et al., 2017), speed of presentation (Tuckute et al., 2024), language (showing generalizability across 45 spoken languages: Malik-Moraleda, Ayyash et al. 2022; Wolna et al., 2024; and sign languages; Richardson et al., 2020; Terhune-Cotter, 2025), materials (Casto et al., 2026; Scott et al., 2017; see also **Supplemental Figure 5**) and aspects of experimental procedure (Lipkin et al., 2022). Importantly, this study reinforces prior findings that a passive language comprehension version of the localizer works well for identifying the language areas (e.g., Fedorenko et al., 2010; Olson et al., 2024; Scott et al., 2017). Of course, in some cases, having a behavioral task—to ensure deep engagement for participants who may struggle with attention, or to directly measure how well the stimuli are understood—may be desirable. Or, one may need to ensure similar levels of behavioral performance when comparing across populations (e.g., Wilson et al., 2017). In such cases, or when creating new versions of a language localizer, it is important to ensure that the contrast is not difficulty-confounded and does not engage parts of the multiple demand (MD) network in addition to the language regions. For example, some past studies have used a contrast between listening to sentences and answering questions about the content vs. listening to sentences played backwards and deciding whether a beep is present at the end (e.g., Newport et al., 2022; Olulade et al., 2020). Such difficulty-confounded contrasts will engage both the language network, but additionally the domain-general Multiple Demand network, which is sensitive to effort across diverse tasks (Duncan et al., 2010; Fedorenko et al., 2013; Shashidhara et al., 2019; Assem et al., 2020)). Such co-engagement of the language and the MD networks for demanding language tasks has been reported across a few language paradigms (e.g., Diachek et al., 2020; Quillen et al., 2021; Wolna et al., 2024). Given that these two networks are functionally distinct, , and the MD network plays a limited role in language processing (Wehbe et al., 2021; Shain et al., 2022; see Fedorenko & Shain, 2022 for a review), conflating them is undesirable and will lead to interpretive challenges.

Finally, it is worth noting that the language network can be identified within individual brains without any functional contrast, from patterns of functional connectivity alone. Braga et al. (2020; see also Du et al., 2024) have shown that this approach works for resting state data; and Du et al. (2025) and Shain & Fedorenko (2025) have extended this result to functional connectivity patterns extracted from random task data. So, those who remain skeptical about functional localizers may prefer to use these task-free data-driven ways of identifying the language areas. However, using localizers remains the most time-efficient approach: a single ∼5 minute run (or even less; Lee et al., 2024) is sufficient in almost all cases.

### 4.3 The MD network is robustly modulated by task demands

Across analyses, the MD network showed the characteristic signature of a domain-general system sensitive to general cognitive effort, distinct from the activation profiles of the language network, consistent with prior findings (Fedorenko et al., 2012, 2013; Hiersche et al., 2024; Mineroff, Blank et al., 2018). In the ***univariate analyses***, responses increased with task difficulty across both sentences and nonwords. Harder tasks, as measured behaviorally (i.e., hard and easy memory probe, comprehension question, sentiment judgment), elicited substantially stronger activation in the MD network than the easier (button-press) tasks. Further, behavioral difficulty (accuracies, response times) reliably predicted MD responses across conditions. These results add to the growing body of evidence that the MD network is robustly engaged during diverse demanding tasks, including classic executive function tasks (Assem et al., 2020; Duncan & Owen, 2000; Fedorenko et al., 2013; Shashidhara et al., 2019) but also tasks in various domains of perception and cognition that may be associated with executive demands. For example, when language processing is accompanied by task demands above and beyond linguistic processing demands related to word recognition or syntactic parsing, the MD network shows above-baseline activity (e.g., Diachek et al., 2020; Quillen et al., 2021; Wolna et al., 2024). Importantly, as noted above, linguistic demands related to the difficulty of accessing words from memory, or building syntactic structures are selectively supported by the language network (see Fedorenko & Shain, 2022 for a review).

Mirroring the univariate results, in the ***multivariate analyses***, the within-task activation patterns were more reliable for more demanding tasks, cross-task pattern similarity reflected task difficulty rather than stimulus type, and task was more robustly decodable from the MD network’s activation patterns than from those in the language network. Although the activations in the MD network allowed for weak decoding of stimulus type, the activations were dominated by task information. These findings align with prior work showing that the MD network can adaptively code both task and stimulus information (Woolgar et al., 2011), and with findings from the visual domain where task information was also shown to be more strongly present in the MD network, whereas stimulus information was more strongly represented in high-level visual regions (Shashidhara et al., 2024). We extend this pattern to language: linguistic stimulus information is primarily represented in the language network, but information on what task is being performed on those stimuli is strongly represented in the MD network.

Taken together, the results reinforce the view that the MD network implements task-general computations that operate independently of the language network. When language processing is embedded within a cognitively demanding paradigm, MD regions become engaged—alongside the language network— to represent task structure and support task-relevant demands, while the language network performs specifically linguistic computations. This division of computational labor explains why language paradigms that include additional task demands systematically recruit two distinct networks and thereby risk conflating language-selective responses with domain-general, task-related responses.

### 4.4 Limitations

Several limitations of the current study suggest clear directions for future work. First,our task battery—although the largest in evaluating effects of task on language network localization—captures only a small fraction of the behaviors that constitute functional linguistic competence (Mahowald, Ivanova et al., 2024), i.e. the use of language in real-life contexts. All six tasks in our study used isolated sentences or simple question-answer pairs. The memory-probe tasks focused on the form of the sentence (asking about particular words comprising it), and the two higher-level tasks (comprehension questions, and sentiment judgments) required deeper engagement with the sentence’s meaning but did not require discourse-level integration, reference resolution, pragmatic inference, domain-specific or abstract reasoning, or planning, which are central components of functional language use. Evaluating language processing in more complex paradigms would tell us whether the stable stimulus-driven signature of the language network persists when linguistic processing is embedded in broader reasoning or communicative contexts.

Second, our sample consists of young, neurotypical adults. As a result, extending these findings to more diverse populations—including children and older adults, non-native speakers, and clinical populations—remains an important future goal.

Finally, fully dissociating stimulus and task processing is challenging: tasks that require additional memory or inference plausibly encourage participants to reprocess or rehearse the sentences, which may lead to additional linguistic computation. As a result, an increase in the response of the language network for more demanding tasks (as we observed) is difficult to interpret unambiguously. Designing experiments that minimize this ambiguity will require some creativity.

## 5 Conclusion

This study demonstrates that the language network exhibits robust, stimulus-driven responses to sentences regardless of task context. Across six task variants—including passive reading, memory probes, comprehension questions, and sentiment judgments—language-selective regions responded consistently more strongly to sentences than to nonwords. Furthermore, the language fROIs defined using any of these tasks were topographically almost as similar as when using different portions of the data from the same task, supporting the reliability and generalizability of functional localization methods for mapping the language network. Although task demands modulated the magnitude of responses within the language network, the activation pattern remained largely stable, suggesting that these regions are consistently engaged in linguistic input processing—demands that were stable across tasks.

These findings clarify the nature of task modulation in the language network: attention or memory demands can amplify responses but do not fundamentally shift the topography of the language-selective regions. Instead, a distinct network—the domain-general MD network—is preferentially engaged by cognitive task demands and strongly encodes task identity, while the language network primarily encodes linguistic content. In this way, the language network appears akin to perceptual brain areas, maintaining stable stimulus representations across task contexts..

## Ethics

The study protocol was approved by MIT’s Committee on the Use of Humans as Experimental Subjects (COUHES), and all participants provided written informed consent in accordance with protocol requirements.

## Data and Code Availability

The experiment scripts, data, codes are available at https://github.com/RuiminGao/DiffTasks.git. The language and the MD parcels are available at https://www.evlab.mit.edu/resources-all/download-parcels. Activation maps of participants are available at https://osf.io/zyfmu.

## Author Contributions

RG: Data curation, Formal analysis, Methodology, Software, Writing - Original Draft, Visualization, Validation; CC: Formal analysis, Methodology, Visualization, Writing - Original Draft; MS: Data curation, Formal analysis, Investigation, Software, Validation; AP: Data curation, Formal analysis, Methodology, Validation; HK: Data curation, Formal analysis, Investigation, Validation; AT: Data curation, Formal analysis, Validation; EF: Conceptualization, Funding acquisition, Methodology, Project administration, Resources, Supervision, Validation, Writing - Review & Editing; AI: Data curation, Investigation, Methodology, Project administration, Supervision, Validation, Writing - Review & Editing.

## Acknowledgements

We acknowledge the Athinoula A. Martinos Imaging Center at the McGovern Institute for Brain Research, MIT, including the technical support team (Steve Shannon and Atsushi Takahashi). We thank EvLab members for assistance with data collection and LIT lab members for feedback on this work. RG and AI were supported by Georgia Tech School of Psychology startup funds. EF was supported by research funds from the McGovern Institute for Brain Research, MIT’s Simons Center for the Social Brain, MIT’s Poitras Center for Psychiatric Disorders Research, and MIT’s Quest for Intelligence.

