## Supplemental materials for "The language network responds robustly to sentences across diverse tasks"

### Supplementary Information

**Behavioral Results**

We found that accuracy decreased for tasks involving higher cognitive demands relative to the button press task. Interactions revealed substantial accuracy reductions relative to the button press task for the nonword condition paired with the hard memory probe ($\beta = -3.89$, $SE = 0.34$, $p < .001$), easy memory probe ($\beta = -0.92$, $SE = 0.45$, $p = .041$), sentiment ($\beta = -2.47$, $SE = 0.57$, $p < .001$), and comprehension question ($\beta = -2.60$, $SE = 0.42$, $p < .001$) tasks. For the sentence condition, accuracy significantly decreased when two sentences were presented compared to one ($\beta = -1.18$, $SE = 0.31$, $p < .001$). A significant interaction effect is found for the hard memory probe task and sentence condition ($\beta = 0.74$, $SE = 0.26$, $p = .005$). Additionally, sentence trials had overall higher accuracy compared to nonwords ($\beta = 0.67$, $SE = 0.27$, $p = .012$).

In line with accuracy findings, response times increased for all tasks compared to the button press task. Significant interactions were found between the nonword condition and the hard memory probe ($\beta = 0.43$, $SE = 0.02$, $p < .001$), easy memory probe ($\beta = 0.26$, $SE = 0.02$, $p < .001$), sentiment question ($\beta = 0.23$, $SE = 0.11$, $p = .043$), and comprehension question ($\beta = 1.30$, $SE = 0.04$, $p < .001$) tasks. For the sentence condition, response time significantly increased when two sentences were presented compared to one ($\beta = 0.08$, $SE = 0.01$, $p < .001$). A significant interaction effect is found for the hard memory probe task and sentence condition ($\beta = 0.03$, $SE = 0.01$, $p = .013$). Sentence trial response times were not significantly higher compared to nonword trials ($\beta = -0.02$, $SE = 0.01$, $p = .135$).

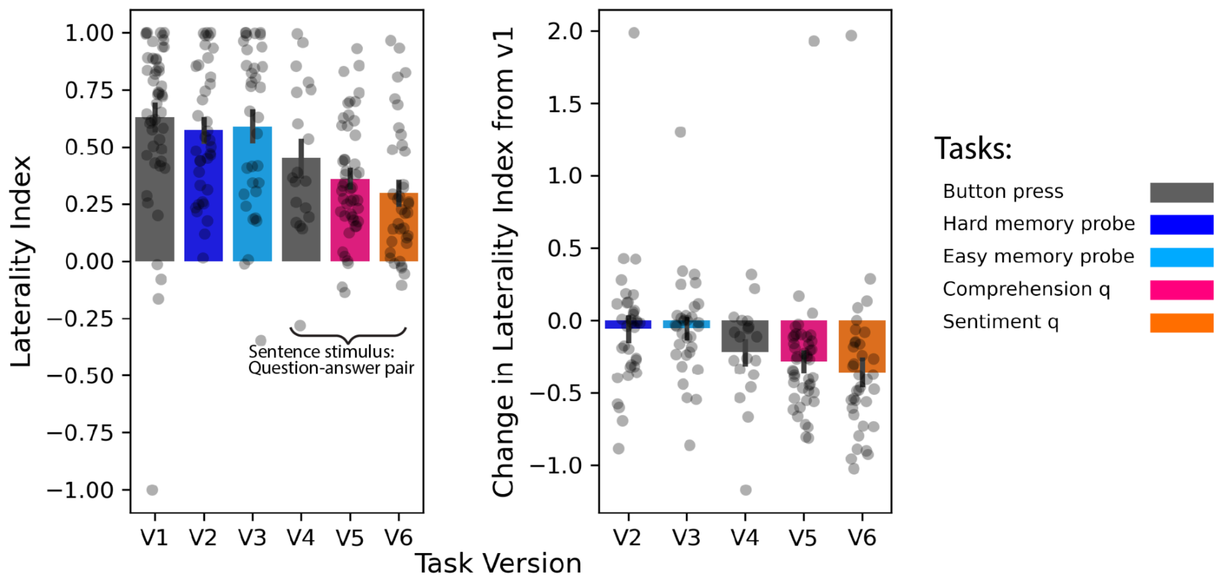


***Supplemental Figure 1.*** *Laterality index across task versions and relative change from V1. Left: Mean laterality index for each task version. Each dot denotes a participant. Right: Change in laterality index for each task version relative to V1. Error bars denote the standard error.*

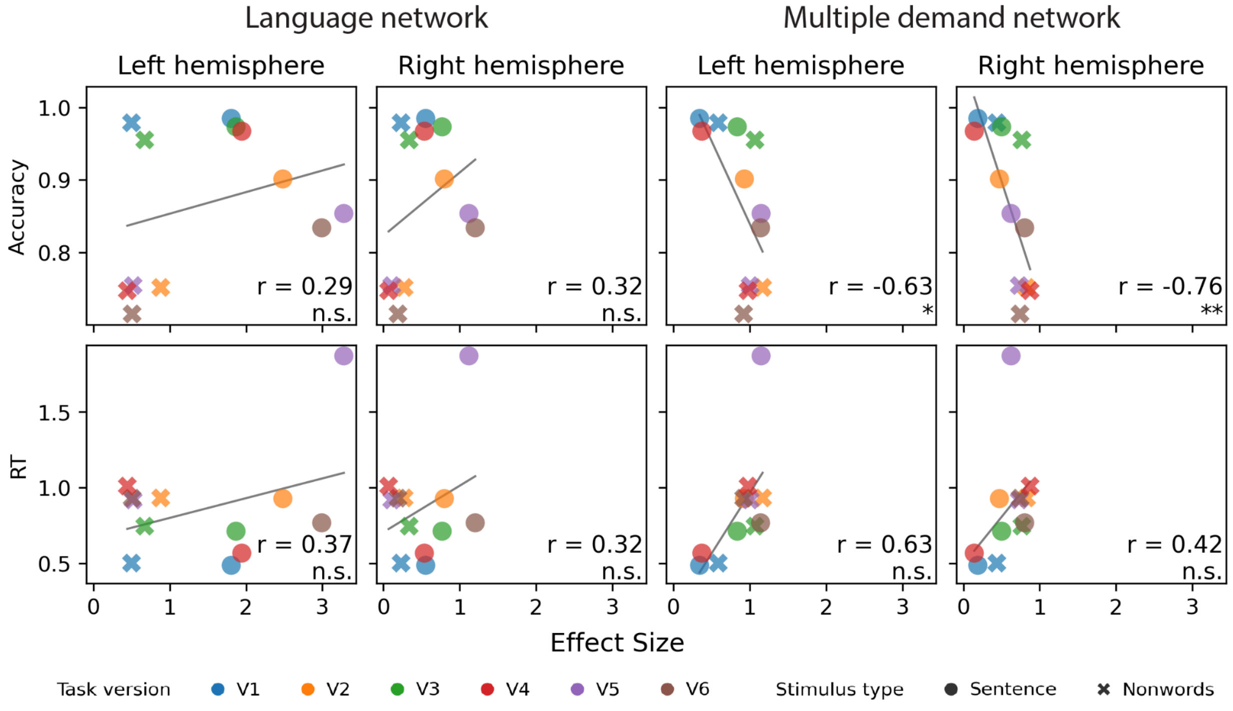


***Supplemental Figure 2.*** *Average accuracy and response time versus the average effect sizes in the language and MD networks of language localizer tasks. FDR correction was conducted across the two hemispheres within each analysis and condition.*

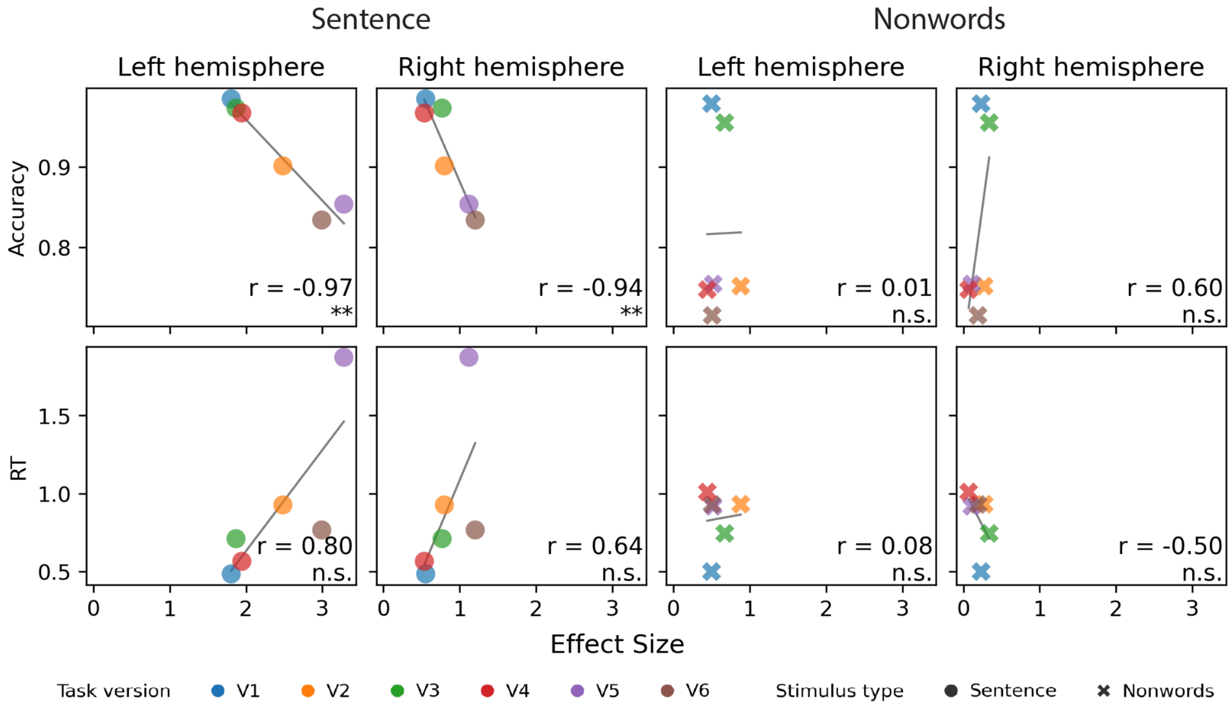


***Supplemental Figure 3.*** *Average accuracy and response time versus the average effect sizes in the language networks of language localizer tasks by stimulus type. FDR correction was conducted across the two hemispheres within each analysis and condition.*

***Supplemental Table 1.*** *Procedure and timing language localizer tasks. Each row denotes event sequences and durations of a trial.*

| Version | Stimulus Type | Initial Blank | Stimulus Presentation | Probe | Post-event Blank | Total Trial Duration |
|---|---|---|---|---|---|---|
| V1 (button press) | Both | 0.1 s | 4.8 s (12 × 0.4 s) | 0.4 s | 0.7 s | 6.0 s |
| V2 (hard memory probe) | Both | 0.1 s | 4.8 s | 1.0 s | 1.0 s | 7.0 s |
| V3 (easy memory probe) | Both | 0.1 s | 4.8 s | 1.0 s | 1.0 s | 7.0 s |
| V4 (button press) | Sentences | 0.1 s | 5.2 s (13 × 0.4 s) | 0.4 s | 0.6 s | 9.0 s |
| V4 (hard memory probe) | Nonwords | 0.1 s | 4.8 s | 2.6 s | 0.4 s | 8.0 s |
| V5 (comprehension q) | Sentences | 0.1 s | 4.8 s | 2.6 s | 0.4 s | 8.0 s |
| V5 (hard memory probe) | Nonwords | 0.1 s | 4.8 s | 2.6 s | 0.4 s | 8.0 s |
| V6 (sentiment q) | Sentences | 0.1 s | 5.2 s (13 × 0.4 s) | 0.4 s | 0.6 s | 9.0 s |
| V6 (hard memory probe) | Nonwords | 0.1 s | 4.8 s | 2.6 s | 0.4 s | 8.0 s |

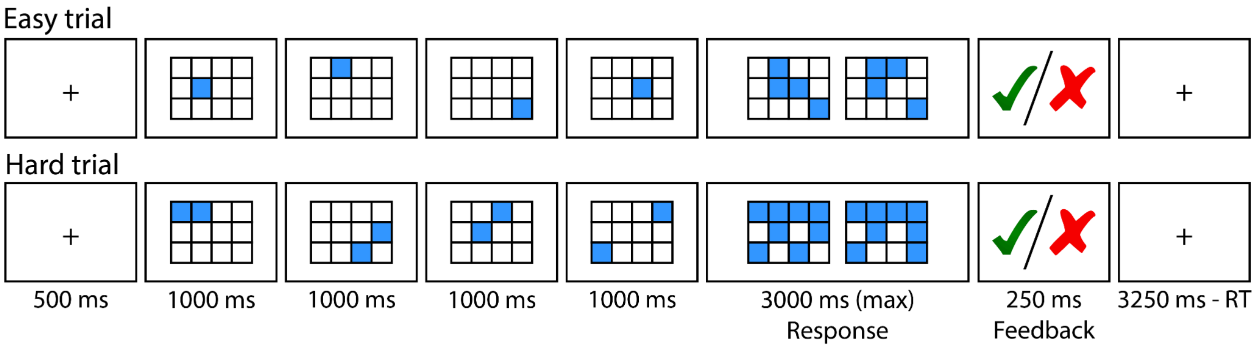


***Supplemental Figure 4.*** *Procedure and timing for the spatial working memory task for localizing MD fROIs.*

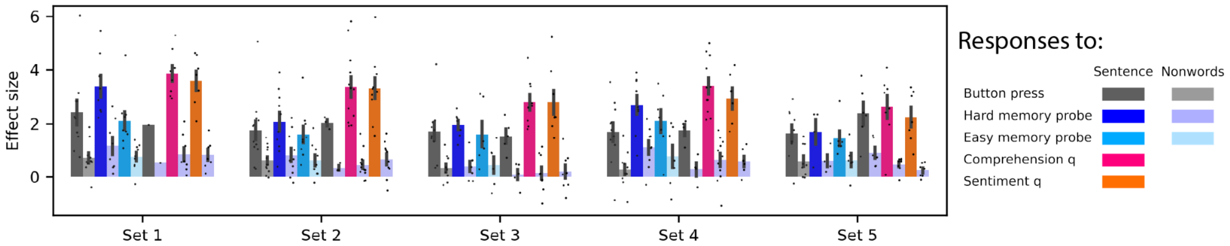


***Supplemental Figure 5.*** *Responses of the core left-hemisphere language network to the sentence and nonword conditions by localizer material set.*

***Supplemental Table 2.*** *Mixed-effect linear regression results for the left language networks on response magnitude by localizer task version. S - sentences, N - nonwords. Sentence (S) condition was used as the reference level (intercept) in the model.*

| Localizer | V1 | | V2 | | V3 | | V4 | | V5 | | V6 | |
|---|---|---|---|---|---|---|---|---|---|---|---|---|
| Regression Term | Beta | Sig | Beta | Sig | Beta | Sig | Beta | Sig | Beta | Sig | Beta | Sig |
| **(Intercept)** | **1.85** | *** | **1.84** | *** | **1.68** | *** | **1.49** | *** | **1.73** | *** | **1.79** | *** |
| *Linguistic demand* | | | | | | | | | | | | |
| QuestionAns>OneSentence | **0.34** | * | **0.43** | * | 0.17 | | **0.23** | * | **0.43** | ** | **0.43** | ** |
| *Task demand* | | | | | | | | | | | | |
| **OneSentence:V2_MemoryAll>V1_ButtonPress** | **0.50** | *** | **0.41** | ** | **0.53** | *** | **0.60** | ** | **0.57** | *** | **0.60** | *** |
| OneSentence:V3_MemoryLast>V1_ButtonPress | 0.01 | | 0.00 | | -0.01 | | -0.32 | | 0.06 | | 0.03 | |
| **QuestionAns:V5_ComprehensionQ>V4_ButtonPress** | **0.85** | *** | **0.88** | ** | **1.00** | *** | **0.85** | ** | **1.04** | *** | **1.05** | *** |
| **QuestionAns:V6_SentimentQ>V4_ButtonPress** | **1.14** | *** | **1.14** | *** | **1.44** | *** | **1.21** | *** | **1.38** | *** | **1.42** | *** |
| *Stimulus type* | | | | | | | | | | | | |
| **OneSentence:V1_ButtonPress:N>S** | **-1.34** | *** | **-1.44** | *** | **-1.28** | *** | **-1.12** | *** | **-1.34** | *** | **-1.28** | *** |
| **OneSentence:V2_MemoryAll:N>S** | **-1.52** | *** | **-1.64** | *** | **-1.55** | *** | **-1.72** | *** | **-1.64** | *** | **-1.75** | *** |
| **OneSentence:V3_MemoryLast:N>S** | **-1.20** | *** | **-1.28** | *** | **-1.16** | *** | **-0.90** | *** | **-1.23** | *** | **-1.12** | *** |
| **QuestionAns:V4_ButtonPress:N>S** | **-1.61** | *** | **-1.87** | *** | **-1.49** | *** | **-1.55** | *** | **-1.70** | *** | **-1.76** | *** |
| **QuestionAns:V5_ComprehensionQ:N>S** | **-2.79** | *** | **-3.04** | *** | **-2.88** | *** | **-2.73** | *** | **-3.17** | *** | **-3.22** | *** |
| **QuestionAns:V6_SentimentQ:N>S** | **-2.52** | *** | **-2.76** | *** | **-2.50** | *** | **-2.45** | *** | **-2.84** | *** | **-2.89** | *** |

***Supplemental Table 3.*** *Mixed-effect linear regression results for the left language networks on response magnitude by fROI. S - sentences, N - nonwords. Sentence (S) condition was used as the reference level (intercept) in the model.*

| ROI | IFGorb | | IFG | | MFG | | AntTemp | | PostTemp | |
|---|---|---|---|---|---|---|---|---|---|---|
| Regression Term | Beta | Sig. | Beta | Sig. | Beta | Sig. | Beta | Sig. | Beta | Sig. |
| **(Intercept)** | **1.64** | *** | **1.87** | *** | **2.54** | *** | **1.17** | *** | **1.73** | *** |
| *Linguistic demand* | | | | | | | | | | |
| QuestionAns>OneSentence | 0.15 | | **0.38** | *** | **0.61** | *** | **0.18** | *** | **0.38** | *** |
| *Task demand* | | | | | | | | | | |
| OneSentence:V2_MemoryAll>V1_ButtonPress | **0.35** | * | **0.87** | *** | **0.81** | *** | 0.08 | | **0.30** | ** |
| OneSentence:V3_MemoryLast>V1_ButtonPress | -0.18 | | 0.21 | | 0.07 | | -0.03 | | -0.07 | |
| **QuestionAns:V5_ComprehensionQ>V4_ButtonPress** | **1.52** | *** | **1.89** | *** | **1.55** | *** | **0.62** | *** | **0.84** | *** |
| **QuestionAns:V6_SentimentQ>V4_ButtonPress** | **1.14** | *** | **1.49** | *** | **1.05** | *** | **0.55** | *** | **0.68** | *** |
| *Stimulus type* | | | | | | | | | | |
| **OneSentence:V1_ButtonPress:N>S** | **-1.22** | *** | **-1.44** | *** | **-1.74** | *** | **-1.03** | *** | **-1.27** | *** |
| **OneSentence:V2_MemoryAll:N>S** | **-1.47** | *** | **-1.98** | *** | **-1.91** | *** | **-1.17** | *** | **-1.50** | *** |
| **OneSentence:V3_MemoryLast:N>S** | **-1.16** | *** | **-1.40** | *** | **-1.43** | *** | **-0.99** | *** | **-1.21** | *** |
| **QuestionAns:V4_ButtonPress:N>S** | **-1.57** | *** | **-1.65** | *** | **-1.97** | *** | **-1.47** | *** | **-1.80** | *** |
| **QuestionAns:V5_ComprehensionQ:N>S** | **-3.20** | *** | **-3.69** | *** | **-3.53** | *** | **-2.03** | *** | **-2.60** | *** |
| **QuestionAns:V6_SentimentQ:N>S** | **-2.79** | *** | **-3.33** | *** | **-3.10** | *** | **-1.96** | *** | **-2.48** | *** |

***Supplemental Table 4.*** *Mixed-effect linear regression results for the left and right language networks on response magnitude. S - sentences, N - nonwords. Sentence (S) condition was used as the reference level (intercept) in the model.*

| Hemisphere | Left hemisphere | | | | Right hemisphere | | | |
|---|---|---|---|---|---|---|---|---|
| Regression Term | Beta | SE | p value | | Beta | SE | p value | |
| **(Intercept)** | **1.79** | **0.27** | **<.001** | *** | **0.57** | **0.11** | **<.001** | *** |
| *Linguistic demand* | | | | | | | | |
| QuestionAns>OneSentence | **0.34** | **0.06** | **<.001** | *** | -0.01 | 0.04 | 0.740 | |
| *Task demand* | | | | | | | | |
| OneSentence:V2_MemoryAll>V1_ButtonPress | **0.48** | **0.09** | **<.001** | *** | 0.06 | 0.07 | 0.387 | |
| OneSentence:V3_MemoryLast>V1_ButtonPress | 0.01 | 0.11 | 0.959 | | 0.13 | 0.10 | 0.189 | |
| **QuestionAns:V5_ComprehensionQ>V4_ButtonPress** | **1.28** | **0.13** | **<.001** | *** | **0.59** | **0.10** | **<.001** | *** |
| **QuestionAns:V6_SentimentQ>V4_ButtonPress** | **0.97** | **0.16** | **<.001** | *** | **0.72** | **0.12** | **<.001** | *** |
| *Stimulus type* | | | | | | | | |
| **OneSentence:V1_ButtonPress:N>S** | **-1.34** | **0.09** | **<.001** | *** | **-0.37** | **0.06** | **<.001** | *** |
| **OneSentence:V2_MemoryAll:N>S** | **-1.61** | **0.09** | **<.001** | *** | **-0.50** | **0.06** | **<.001** | *** |
| **OneSentence:V3_MemoryLast:N>S** | **-1.24** | **0.09** | **<.001** | *** | **-0.48** | **0.06** | **<.001** | *** |
| **QuestionAns:V4_ButtonPress:N>S** | **-1.69** | **0.14** | **<.001** | *** | **-0.52** | **0.10** | **<.001** | *** |
| **QuestionAns:V5_ComprehensionQ:N>S** | **-3.00** | **0.12** | **<.001** | *** | **-1.14** | **0.09** | **<.001** | *** |
| **QuestionAns:V6_SentimentQ:N>S** | **-2.72** | **0.16** | **<.001** | *** | **-1.21** | **0.13** | **<.001** | *** |

***Supplemental Table 5.*** *Mixed-effect linear regression results for the left and right MD networks on response magnitude. S - sentences, N - nonwords. Sentence (S) condition was used as the reference level (intercept) in the model.*

| Hemisphere | Left hemisphere | | | | Right hemisphere | | | |
|---|---|---|---|---|---|---|---|---|
| Regression Term | Beta | SE | p value | | Beta | SE | p value | |
| (Intercept) | **0.37** | **0.15** | **0.025** | * | 0.20 | 0.12 | 0.113 | |
| *Linguistic demand* | | | | | | | | |
| QuestionAns>OneSentence | 0.05 | 0.07 | 0.421 | | **-0.19** | **0.06** | **0.002** | ** |
| *Task demand* | | | | | | | | |
| **OneSentence:V2_MemoryAll>V1_ButtonPress** | **0.51** | **0.09** | **<.001** | *** | **0.28** | **0.11** | **0.013** | * |
| **OneSentence:V3_MemoryLast>V1_ButtonPress** | **0.45** | **0.10** | **<.001** | *** | **0.38** | **0.11** | **0.002** | ** |
| **QuestionAns:V5_ComprehensionQ>V4_ButtonPress** | **0.72** | **0.10** | **<.001** | *** | **0.59** | **0.12** | **<.001** | *** |
| **QuestionAns:V6_SentimentQ>V4_ButtonPress** | **0.77** | **0.13** | **<.001** | *** | **0.85** | **0.16** | **<.001** | *** |
| *Stimulus type* | | | | | | | | |
| **OneSentence:V1_ButtonPress:N>S** | **0.23** | **0.07** | **0.001** | ** | **0.24** | **0.07** | **0.001** | ** |
| **OneSentence:V2_MemoryAll:N>S** | **0.22** | **0.08** | **0.005** | ** | **0.31** | **0.08** | **<.001** | *** |
| **OneSentence:V3_MemoryLast:N>S** | **0.18** | **0.08** | **0.021** | * | **0.19** | **0.08** | **0.021** | * |
| **QuestionAns:V4_ButtonPress:N>S** | **0.73** | **0.13** | **<.001** | *** | **0.90** | **0.13** | **<.001** | *** |
| QuestionAns:V5_ComprehensionQ:N>S | -0.13 | 0.08 | 0.121 | | 0.11 | 0.09 | 0.186 | |
| QuestionAns:V6_SentimentQ:N>S | -0.16 | 0.10 | 0.122 | | -0.02 | 0.12 | 0.853 | |

***Supplemental Table 6.*** *Mixed-effect linear regression results for the left and right DMN networks on response magnitude. S - sentences, N - nonwords. Sentence (S) condition was used as the reference level (intercept) in the model.*

| Hemisphere | Left hemisphere | | | | Right hemisphere | | | |
|---|---|---|---|---|---|---|---|---|
| Regression Term | Beta | SE | p value | | Beta | SE | p value | |
| (Intercept) | 0.08 | 0.13 | 0.530 | | -0.13 | 0.09 | 0.175 | |
| *Linguistic demand* | | | | | | | | |
| QuestionAns>OneSentence | -0.01 | 0.08 | 0.852 | | -0.01 | 0.07 | 0.906 | |
| *Task demand* | | | | | | | | |
| **OneSentence:V2_MemoryAll>V1_ButtonPress** | **-0.33** | **0.08** | **<.001** | *** | **-0.36** | **0.08** | **<.001** | *** |
| OneSentence:V3_MemoryLast>V1_ButtonPress | -0.07 | 0.09 | 0.419 | | 0.02 | 0.08 | 0.845 | |
| **QuestionAns:V5_ComprehensionQ>V4_ButtonPress** | **-0.30** | **0.10** | **0.003** | ** | **-0.39** | **0.09** | **<.001** | *** |
| QuestionAns:V6_SentimentQ>V4_ButtonPress | -0.25 | 0.14 | 0.076 | | **-0.35** | **0.13** | **0.008** | ** |
| *Stimulus type* | | | | | | | | |
| OneSentence:V1_ButtonPress:N>S | 0.05 | 0.07 | 0.466 | | **0.15** | **0.06** | **0.017** | * |
| OneSentence:V2_MemoryAll:N>S | -0.02 | 0.08 | 0.802 | | 0.13 | 0.07 | 0.072 | |
| OneSentence:V3_MemoryLast:N>S | -0.06 | 0.08 | 0.436 | | 0.01 | 0.07 | 0.859 | |
| **QuestionAns:V4_ButtonPress:N>S** | **-0.51** | **0.13** | **<.001** | *** | **-0.32** | **0.12** | **0.006** | ** |
| QuestionAns:V5_ComprehensionQ:N>S | -0.09 | 0.08 | 0.280 | | 0.13 | 0.07 | 0.057 | |
| QuestionAns:V6_SentimentQ:N>S | -0.11 | 0.11 | 0.306 | | 0.15 | 0.08 | 0.080 | |

***Supplemental Table 7.*** *Mixed-effect linear regression results for the left and right language networks on spatial correlation. S - sentences, N - nonwords.*

| Hemisphere | Left hemisphere | | | | Right hemisphere | | | |
|---|---|---|---|---|---|---|---|---|
| Regression Term | Beta | SE | p value | | Beta | SE | p value | |
| **(Intercept)** | **0.53** | **0.11** | **0.002** | ** | **0.31** | **0.07** | **0.003** | ** |
| *Stimulus type* | | | | | | | | |
| N.S>N.N | -0.01 | 0.01 | 0.302 | | -0.01 | 0.01 | 0.531 | |
| **S.S>N.N** | **0.36** | **0.01** | **<.001** | *** | **0.15** | **0.01** | **<.001** | *** |
| *Linguistic demand* | | | | | | | | |
| **N.S:OneSentence.QuestionAns>OneSentence.OneSentence** | **-0.08** | **0.00** | **<.001** | *** | **-0.02** | **0.00** | **<.001** | *** |
| S.S:OneSentence.QuestionAns>OneSentence.OneSentence | **-0.02** | **0.00** | **<.001** | *** | 0.01 | 0.00 | 0.280 | |
| **S.S:QuestionAns.QuestionAns>OneSentence.OneSentence** | **0.19** | **0.01** | **<.001** | *** | **0.14** | **0.01** | **<.001** | *** |
| *Task demand* | | | | | | | | |
| N.N:ButtonPress.Task>ButtonPress.ButtonPress | 0.02 | 0.01 | 0.175 | | 0.01 | 0.01 | 0.445 | |
| **N.N:Task.Task>ButtonPress.ButtonPress** | **0.18** | **0.01** | **<.001** | *** | **0.16** | **0.01** | **<.001** | *** |
| N.S:ButtonPress.Task>ButtonPress.ButtonPress | 0.01 | 0.01 | 0.188 | | 0.00 | 0.01 | 0.846 | |
| **N.S:Task.Task>ButtonPress.ButtonPress** | **0.07** | **0.01** | **<.001** | *** | **0.08** | **0.01** | **<.001** | *** |
| S.S:ButtonPress.Task>ButtonPress.ButtonPress | -0.01 | 0.01 | 0.218 | | **0.03** | **0.01** | **<.001** | *** |
| **S.S:Task.Task>ButtonPress.ButtonPress** | **0.14** | **0.01** | **<.001** | *** | **0.16** | **0.01** | **<.001** | *** |

***Supplemental Table 8.*** *Mixed-effect linear regression results for the left and right MD networks on spatial correlation. S - sentences, N - nonwords.*

| Hemisphere | Left hemisphere | | | | Right hemisphere | | | |
|---|---|---|---|---|---|---|---|---|
| Regression Term | Beta | SE | p value | | Beta | SE | p value | |
| **(Intercept)** | **0.40** | **0.07** | **<.001** | *** | **0.34** | **0.05** | **<.001** | *** |
| *Stimulus type* | | | | | | | | |
| N.S>N.N | -0.01 | 0.03 | 0.757 | | -0.02 | 0.03 | 0.470 | |
| S.S>N.N | **0.11** | **0.03** | **<.001** | *** | 0.02 | 0.03 | 0.576 | |
| *Linguistic demand* | | | | | | | | |
| **N.S:OneSentence.QuestionAns>OneSentence.OneSentence** | **0.02** | **0.01** | **<.001** | *** | **0.03** | **0.01** | **<.001** | *** |
| S.S:OneSentence.QuestionAns>OneSentence.OneSentence | **0.02** | **0.01** | **0.041** | * | -0.01 | 0.01 | 0.314 | |
| **S.S:QuestionAns.QuestionAns>OneSentence.OneSentence** | **0.18** | **0.01** | **<.001** | *** | **0.11** | **0.01** | **<.001** | *** |
| *Task demand* | | | | | | | | |
| **N.N:ButtonPress.Task>ButtonPress.ButtonPress** | **0.08** | **0.02** | **0.001** | ** | **0.05** | **0.03** | **0.035** | * |
| **N.N:Task.Task>ButtonPress.ButtonPress** | **0.34** | **0.02** | **<.001** | *** | **0.29** | **0.03** | **<.001** | *** |
| N.S:ButtonPress.Task>ButtonPress.ButtonPress | **0.05** | **0.01** | **0.001** | *** | 0.01 | 0.01 | 0.338 | |
| **N.S:Task.Task>ButtonPress.ButtonPress** | **0.26** | **0.01** | **<.001** | *** | **0.22** | **0.01** | **<.001** | *** |
| S.S:ButtonPress.Task>ButtonPress.ButtonPress | 0.02 | 0.01 | 0.173 | | **0.03** | **0.02** | **0.025** | * |
| **S.S:Task.Task>ButtonPress.ButtonPress** | **0.25** | **0.02** | **<.001** | *** | **0.26** | **0.02** | **<.001** | *** |

***Supplemental Table 9.*** *Mixed-effect linear regression results for the left and right DMN networks on spatial correlation. S - sentences, N - nonwords.*

| Hemisphere | Left hemisphere | | | | Right hemisphere | | | |
|---|---|---|---|---|---|---|---|---|
| Regression Term | Beta | SE | p value | | Beta | SE | p value | |
| **(Intercept)** | **0.22** | **0.05** | **<.001** | *** | **0.23** | **0.04** | **<.001** | *** |
| *Stimulus type* | | | | | | | | |
| N.S>N.N | -0.04 | 0.04 | 0.365 | | -0.06 | 0.04 | 0.107 | |
| **S.S>N.N** | **0.14** | **0.04** | **0.001** | *** | **0.10** | **0.04** | **0.015** | * |
| *Linguistic demand* | | | | | | | | |
| N.S:OneSentence.QuestionAns>OneSentence.OneSentence | 0.00 | 0.01 | 0.670 | | 0.01 | 0.01 | 0.246 | |
| S.S:OneSentence.QuestionAns>OneSentence.OneSentence | **0.04** | **0.01** | **0.004** | ** | -0.01 | 0.01 | 0.709 | |
| **S.S:QuestionAns.QuestionAns>OneSentence.OneSentence** | **0.18** | **0.02** | **<.001** | *** | **0.12** | **0.02** | **<.001** | *** |
| *Task demand* | | | | | | | | |
| N.N:ButtonPress.Task>ButtonPress.ButtonPress | -0.03 | 0.04 | 0.330 | | -0.03 | 0.04 | 0.336 | |
| **N.N:Task.Task>ButtonPress.ButtonPress** | **0.13** | **0.04** | **<.001** | *** | **0.14** | **0.04** | **<.001** | *** |
| N.S:ButtonPress.Task>ButtonPress.ButtonPress | -0.02 | 0.02 | 0.334 | | -0.03 | 0.04 | 0.336 | |
| **N.S:Task.Task>ButtonPress.ButtonPress** | **0.07** | **0.02** | **<.001** | *** | **0.14** | **0.04** | **<.001** | *** |
| S.S:ButtonPress.Task>ButtonPress.ButtonPress | 0.02 | 0.02 | 0.283 | | -0.01 | 0.01 | 0.709 | |
| **S.S:Task.Task>ButtonPress.ButtonPress** | **0.15** | **0.02** | **<.001** | *** | **0.12** | **0.02** | **<.001** | *** |

***Supplemental Table 10.*** *Results of post-hoc linear mixed-effects regression tests on fROI overlap in the left and right language networks.*

| Hemisphere | Left hemisphere | | | | Right hemisphere | | | |
|---|---|---|---|---|---|---|---|---|
| Contrast | Beta | SE | p value | | Beta | SE | p value | |
| *Within-participant.Same-task vs Within-participant.Different-task* | | | | | | | | |
| V1 | **0.06** | **0.01** | **<.001** | *** | -0.01 | 0.01 | 1.000 | |
| **V2** | **0.06** | **0.01** | **<.001** | *** | **0.03** | **0.01** | **<.001** | *** |
| V3 | 0.02 | 0.01 | 0.117 | | -0.01 | 0.01 | 1.000 | |
| **V4** | **0.07** | **0.01** | **<.001** | *** | **0.09** | **0.01** | **<.001** | *** |
| **V5** | **0.16** | **0.01** | **<.001** | *** | **0.18** | **0.01** | **<.001** | *** |
| **V6** | **0.13** | **0.01** | **<.001** | *** | **0.16** | **0.01** | **<.001** | *** |
| *Within-participant.Different-task vs Between-participant.Same-task* | | | | | | | | |
| **V1** | **0.33** | **0.01** | **<.001** | *** | **0.19** | **0.01** | **<.001** | *** |
| **V2** | **0.39** | **0.01** | **<.001** | *** | **0.22** | **0.01** | **<.001** | *** |
| **V3** | **0.36** | **0.01** | **<.001** | *** | **0.21** | **0.01** | **<.001** | *** |
| **V4** | **0.38** | **0.01** | **<.001** | *** | **0.22** | **0.01** | **<.001** | *** |
| **V5** | **0.40** | **0.01** | **<.001** | *** | **0.23** | **0.01** | **<.001** | *** |
| **V6** | **0.39** | **0.01** | **<.001** | *** | **0.24** | **0.01** | **<.001** | *** |

***Supplemental Table 11.*** *Results of post-hoc linear mixed-effects regression tests on fROI overlap by ROI in the left language network.*

| ROI | IFGorb | | IFG | | MFG | | AntTemp | | PostTemp | |
|---|---|---|---|---|---|---|---|---|---|---|
| Contrast | Beta | Sig. | Beta | Sig. | Beta | Sig. | Beta | Sig. | Beta | Sig. |
| *Within-participant.Same-task vs Within-participant.Different-task* | | | | | | | | | | |
| **V1** | **0.07** | *** | **0.07** | *** | **0.07** | *** | **0.02** | * | **0.04** | *** |
| **V2** | **0.07** | *** | **0.08** | *** | **0.08** | *** | **0.04** | *** | **0.05** | *** |
| V3 | -0.03 | | **0.05** | * | 0.03 | | 0.01 | | **0.03** | * |
| **V4** | **0.10** | *** | **0.06** | ** | **0.08** | *** | **0.06** | *** | **0.06** | *** |
| **V5** | **0.19** | *** | **0.15** | *** | **0.18** | *** | **0.14** | *** | **0.14** | *** |
| **V6** | **0.12** | *** | **0.13** | *** | **0.16** | *** | **0.11** | *** | **0.13** | *** |
| *Within-participant.Different-task vs Between-participant.Same-task* | | | | | | | | | | |
| **V1** | **0.27** | *** | **0.33** | *** | **0.34** | *** | **0.32** | *** | **0.41** | *** |
| **V2** | **0.30** | *** | **0.40** | *** | **0.40** | *** | **0.36** | *** | **0.47** | *** |
| **V3** | **0.28** | *** | **0.38** | *** | **0.36** | *** | **0.35** | *** | **0.43** | *** |
| **V4** | **0.30** | *** | **0.38** | *** | **0.40** | *** | **0.34** | *** | **0.46** | *** |
| **V5** | **0.33** | *** | **0.41** | *** | **0.41** | *** | **0.38** | *** | **0.46** | *** |
| **V6** | **0.31** | *** | **0.41** | *** | **0.41** | *** | **0.37** | *** | **0.46** | *** |

**Supplemental Table 12.** Multivariate pattern analysis decoding accuracies. SVM: Support Vector Machine, kNN: k-Nearest Neighbors, LR: Logistic Regression. Bonferroni correction for multiple comparisons within each network and classification task.

| Network | Testing Condition | Correlation | | SVM | | kNN | | LR | |
|---|---|---|---|---|---|---|---|---|---|
| | | Accuracy | Sig. | Accuracy | Sig. | Accuracy | Sig. | Accuracy | Sig. |
| *Within-condition training: stimulus type classification* | | | | | | | | | |
| Language (LH) | Button press | 0.67 | *** | 0.73 | *** | 0.66 | *** | 0.69 | *** |
| | hard memory probe | 0.70 | *** | 0.77 | *** | 0.68 | *** | 0.72 | *** |
| | Easy memory probe | 0.69 | *** | 0.73 | *** | 0.65 | *** | 0.68 | *** |
| Language (RH) | Button press | 0.61 | *** | 0.64 | *** | 0.60 | *** | 0.63 | *** |
| | hard memory probe | 0.66 | *** | 0.68 | *** | 0.64 | *** | 0.66 | *** |
| | Easy memory probe | 0.59 | *** | 0.62 | *** | 0.59 | *** | 0.62 | *** |
| MD | Button press | 0.55 | ** | 0.57 | *** | 0.56 | *** | 0.59 | *** |
| | hard memory probe | 0.58 | *** | 0.60 | *** | 0.57 | *** | 0.65 | *** |
| | Easy memory probe | 0.56 | *** | 0.56 | *** | 0.54 | ** | 0.60 | *** |
| *Within-condition training: task classification* | | | | | | | | | |
| Language (LH) | Sentence | 0.46 | *** | 0.51 | *** | 0.41 | *** | 0.48 | *** |
| | Nonwords | 0.39 | *** | 0.48 | *** | 0.38 | ** | 0.42 | *** |
| Language (RH) | Sentence | 0.43 | *** | 0.49 | *** | 0.42 | *** | 0.45 | *** |
| | Nonwords | 0.37 | ** | 0.45 | *** | 0.39 | ** | 0.42 | *** |
| MD | Sentence | 0.44 | *** | 0.52 | *** | 0.44 | *** | 0.56 | *** |
| | Nonwords | 0.42 | *** | 0.52 | *** | 0.40 | *** | 0.53 | *** |
| *Across-condition training: stimulus type classification* | | | | | | | | | |
| Language (LH) | Button press | 0.66 | *** | 0.70 | *** | 0.65 | *** | 0.67 | *** |
| | hard memory probe | 0.70 | *** | 0.75 | *** | 0.65 | *** | 0.69 | *** |
| | Easy memory probe | 0.69 | *** | 0.70 | *** | 0.65 | *** | 0.68 | *** |
| Language (RH) | Button press | 0.62 | *** | 0.65 | *** | 0.63 | *** | 0.64 | *** |
| | hard memory probe | 0.66 | *** | 0.67 | *** | 0.61 | *** | 0.63 | *** |
| | Easy memory probe | 0.61 | *** | 0.62 | *** | 0.59 | *** | 0.58 | *** |
| MD | Button press | 0.56 | ** | 0.58 | *** | 0.56 | *** | 0.60 | *** |
| | hard memory probe | 0.58 | *** | 0.59 | *** | 0.56 | ** | 0.63 | *** |
| | Easy memory probe | 0.56 | ** | 0.56 | *** | 0.56 | *** | 0.58 | *** |
| *Across-condition training: task classification* | | | | | | | | | |
| Language (LH) | Sentence | 0.38 | ** | 0.44 | *** | 0.36 | * | 0.38 | *** |
| | Nonwords | 0.38 | ** | 0.43 | *** | 0.37 | ** | 0.40 | *** |
| Language (RH) | Sentence | 0.39 | *** | 0.45 | *** | 0.38 | ** | 0.41 | *** |
| | Nonwords | 0.39 | *** | 0.45 | *** | 0.39 | *** | 0.41 | *** |
| MD | Sentence | 0.41 | *** | 0.51 | *** | 0.39 | *** | 0.53 | *** |
| | Nonwords | 0.42 | *** | 0.49 | *** | 0.41 | *** | 0.54 | *** |

**Supplemental Table 13.** Binomial mixed-effects model predicting decoding accuracy. Fixed effects included training type, system (LH language vs. MD), testing group, and their interactions. Random intercepts were included for participant and classifier type.

| Regression Term | Estimate | SE | *z* | p value | 95% CI |
|---|---|---|---|---|---|
| *Stimulus type decoding* | | | | | |
| (Intercept) | -0.06 | 0.11 | -0.59 | 0.558 | [-0.27, 0.15] |
| Language_LH:AcrossCondition:TestOnN | -0.39 | 0.05 | -8.60 | <.001 | [-0.48, -0.30] |
| Language_LH:WithinCondition:TestOnN | -0.26 | 0.05 | -5.77 | <.001 | [-0.35, -0.17] |
| Language_LH:AcrossCondition:TestOnS | -0.39 | 0.05 | -8.58 | <.001 | [-0.48, -0.30] |
| Language_LH:WithinCondition:TestOnS | -0.09 | 0.05 | -2.10 | 0.036 | [-0.18, -0.01] |
| MD:AcrossCondition:TestOnN | -0.10 | 0.05 | -2.15 | 0.032 | [-0.19, -0.01] |
| MD:WithinCondition:TestOnN | -0.05 | 0.05 | -1.17 | 0.240 | [-0.14, 0.04] |
| MD:AcrossCondition:TestOnS | -0.10 | 0.05 | -2.30 | 0.021 | [-0.19, -0.02] |
| *Task decoding* | | | | | |
| (Intercept) | 0.27 | 0.08 | 3.43 | <.001 | [0.11, 0.42] |
| Language_LH:AcrossCondition:TestOnV1 | 0.46 | 0.06 | 8.06 | <.001 | [0.35, 0.57] |
| Language_LH:WithinCondition:TestOnV1 | 0.53 | 0.06 | 9.25 | <.001 | [0.42, 0.64] |
| Language_LH:AcrossCondition:TestOnV2 | 0.58 | 0.06 | 10.10 | <.001 | [0.47, 0.69] |
| Language_LH:WithinCondition:TestOnV2 | 0.68 | 0.06 | 11.62 | <.001 | [0.56, 0.79] |
| Language_LH:AcrossCondition:TestOnV3 | 0.50 | 0.06 | 8.82 | <.001 | [0.39, 0.62] |
| Language_LH:WithinCondition:TestOnV3 | 0.56 | 0.06 | 9.66 | <.001 | [0.44, 0.67] |
| MD:AcrossCondition:TestOnV1 | 0.03 | 0.06 | 0.55 | 0.582 | [-0.08, 0.14] |
| MD:WithinCondition:TestOnV1 | 0.01 | 0.06 | 0.13 | 0.896 | [-0.10, 0.12] |
| MD:AcrossCondition:TestOnV2 | 0.10 | 0.06 | 1.86 | 0.062 | [-0.01, 0.21] |
| MD:WithinCondition:TestOnV2 | 0.14 | 0.06 | 2.54 | 0.011 | [0.03, 0.25] |
| MD:AcrossCondition:TestOnV3 | 0.01 | 0.06 | 0.14 | 0.890 | [-0.10, 0.12] |
